# Ergosterol mediates aggregation of natamycin in the yeast plasma membrane

**DOI:** 10.1101/2023.12.05.570146

**Authors:** Maria Szomek, Vibeke Akkerman, Line Lauritsen, Hanna-Loisa Walther, Alice Dupont Juhl, Katja Thaysen, Jacob Marcus Egebjerg, Douglas F. Covey, Max Lehmann, Pablo Wessig, Alexander J. Foster, Bert Poolman, Stephan Werner, Gerd Schneider, Peter Müller, Daniel Wüstner

## Abstract

Polyene macrolides are antifungal substances, which interact with cells in a sterol-dependent manner. While being widely used, their mode of action is poorly understood. Here, we employ ultraviolet-sensitive (UV) microscopy to show that the antifungal polyene natamycin binds to the yeast plasma membrane (PM) and causes permeation of propidium iodide into cells. Right before membrane permeability becomes compromised, we observed clustering of natamycin in the PM that was independent of PM protein domains. Aggregation of natamycin was paralleled by cell deformation and membrane blebbing as revealed by soft X-ray microscopy. Substituting ergosterol for cholesterol decreased natamycin binding and resulted in reduced clustering of natamycin in the PM. Blocking of ergosterol synthesis necessitates sterol import via the ABC transporters Aus1/Pdr11 to ensure natamycin binding. Quantitative imaging of dehydroergosterol (DHE) and cholestatrienol (CTL), two analogs of ergosterol and cholesterol, respectively, revealed a largely homogeneous lateral sterol distribution in the PM, ruling out that natamycin binds to pre-assembled sterol domains. Depletion of sphingolipids using myriocin increased natamycin binding to yeast cells, likely by increasing the ergosterol fraction in the outer PM leaflet. We conclude that ergosterol-specific aggregation of natamycin in the yeast PM underlies its antifungal activity, which can be synergistically enhanced by inhibitors of sphingolipid synthesis.

**Significance:** Ergosterol is the major sterol in the membranes of fungi and a major target for antifungal treatments. Polyene macrolides, such as natamycin, are known to target ergosterol but the underlying mechanisms for their preference for this yeast sterol compared to mammalian cholesterol is not understood. This study shows that natamycin forms aggregates when associated with yeast S. cerevisiae in an ergosterol-dependent manner. Cholesterol can only partially substitute for ergosterol with respect to natamycin binding and aggregation. Membrane-associated aggregation of natamycin is not the result of pre-formed sterol domains in the cell membrane, as we show by direct visualization of minimally modified ergosterol and cholesterol analogs. Inhibiting sphingolipid synthesis increased membrane association and antifungal activity of natamycin, suggesting that targeting sphingolipids in combination with polyene macrolides could lead to novel drug treatment approaches against fungal infections.

## Introduction

Yeast cells have a stringent lateral organization of their plasma membrane (PM) into co-existing domains, defined by distinct protein markers ^1^. This structural hierarchy of protein distribution ensures the highly regulated function of membrane proteins which has to adapt to a variety of environmental conditions. Eisosomes, colocalizing with the membrane compartment containing Can1 (MCC), are sites of surface invaginations, in which membrane proteins such as the MCC marker Sur7 accumulate in a reversible manner ^2–4^. The structure and function of MCC/eisosomes are regulated by membrane tension and thereby directly responsive to changes in environmental conditions ^5^. Under optimal growth conditions MCC/eisosomes form ca. 100-nm sized patches with slight indentation ^6^, but glucose starvation or hyperosmotic conditions, which both cause cell shrinkage, can result in deepened eisosomes, in which transporters become trapped ^5^. Vice versa, hypoosmotic conditions result in flattening of eisosomes and export of membrane proteins from this compartment ^5^. Sequestration of membrane transporters in eisosomes is often regulated by their transport cycle, as addition of substrates, for example to the methionine transporter Mup1 or the lysine transporter Lyp1, causes export from this compartment ^5,7^. Another prominent PM domain is the membrane compartment containing Pma1 (MCP), which is supposed to be enriched in certain transporters including the proton pump Pma1 as well as in sphingolipids ^8^. In contrast to this extensive knowledge about protein dynamics and compartmentalization in the yeast PM, our understanding of the distribution of lipids including sterols is very limited. Ergosterol, the main sterol in yeast, has been suggested to co-cluster with eisosome markers, but these conclusions were only drawn from the partitioning of a polyene macrolide, filipin, which could self-aggregate in the membrane ^2,9^. In a recent study, using lipidomics of isolated PM domains, we found comparable ergosterol abundance in MCC and MCP, suggesting that the lateral distribution of ergosterol in the PM is rather homogeneous ^8^. In addition to the lateral sterol organization, the transbilayer distribution of ergosterol in the yeast PM is important. By replacing ergosterol in yeast with an intrinsically fluorescent ergosterol analogue, dehydroergosterol (DHE), which differs from ergosterol only in having one additional double bond, we found a highly asymmetric sterol distribution in the yeast PM with the majority of DHE in the inner PM leaflet ^10^. Others observed that yeast sphingolipids, which are characterized by long and saturated acyl chains, form gel-like domains in the PM, which are largely ergosterol free ^11^. Thus, an attractive model is that sphingolipids with their large hydrophilic head groups reside primarily in the outer leaflet, while ergosterol locates mostly in the inner PM leaflet in the unperturbed yeast PM. Supporting that model, we found that myriocin, which blocks sphingolipid synthesis caused a reallocation of DHE to the outer PM leaflet ^10^.

Polyene macrolides are natural antifungal compounds, which are not only used to monitor cellular sterol pools, as shown for filipin, but also to treat fungal infections, as known from amphotericin B (AmpB), nystatin or natamycin. These polyenes target ergosterol in the yeast PM, but their precise mode of action is often poorly understood. While AmpB and nystatin form ion pores in the membrane, natamycin kills yeast by a different but poorly defined mechanism ^12,13^. Natamycin is often used to treat fungal keratitis but also to protect food products against fungal molds ^14^. Treating fungi with natamycin inhibits nutrient uptake by secondary active transporters and causes rapid upregulation of their expression but does not result in membrane pore formation ^13,15,16^. We showed recently, that natamycin forms oligomers in model membranes, where it interferes with the ergosterol-dependent activity of the reconstituted lysine transporter Lyp1 ^17^. We also found that natamycin immobilizes ergosterol and cholesterol to a comparable extent but interferes preferentially with the ergosterol-induced liquid-ordered phase in model membranes ^17,18^. How these properties of natamycin relate to its action on the yeast PM is unknown.

A common property of polyene macrolides is their amphiphilic character due to their polar sugar moiety and polyphenol region on one side and the apolar polyene structure on the other part of the molecules. Due to these structural features, polyenes form aggregates in aqueous solution above a critical aggregation concentration (CAC) which has been determined not only for nystatin (CAC=3 µM) and AmpB (CAC=1.1 µM) but also for natamycin by various methods (CAC=20.9-82.5 µM) ^14,17,19–21^. Whether self-aggregation of polyenes is required for their antifungal activity is debated. Some studies show that the hemolytic activity of larger polyenes, like AmpB but also of tetraenes, like lucensomycin, correlates with their respective CAC ^20,22,23^. On the other hand, increasing the solubility of polyenes, for example, by incorporating them into liposomes or by complexation with albumin or cyclodextrin decreases their toxicity to mammalian cells without interfering with their antifungal activity ^20,21,24,25^. These and similar results indicate that extracellular aggregates are toxic to mammalian cells but not to yeast ^26,27^, putting the aggregation behavior of polyenes on the center stage of their antifungal selectivity. Recent studies by Burke and co-workers put forward an alternative model, according to which extracellular aggregates of AmpB, nystatin, and natamycin would extract ergosterol from yeast cells, suggesting that such aggregates (called ‘polyene sponges’) are directly responsible for the antifungal activity of glycosylated polyenes macrolides ^12,28,29^. However, at least for nystatin and natamycin, the evidence for such a model is circumstantial as it is based on the observation that pre-complexation of these polyenes with sterols reduces their antifungal activity ^12^. Mixing polyenes with sterols in water indeed causes formation of sterol-polyene complexes ^17,23,29,30^, but this could also interfere with initial polyene binding to the yeast membrane as well as with sterol extraction into extracellular polyene sponges. AmpB and nystatin rapidly bind to model membranes, and we have recently shown that this is also the case for natamycin ^17^. Initial binding is followed by slower membrane rearrangements including self-aggregation of the polyenes in the bilayer. Nystatin and AmpB assemble into ion pores in the membrane, while smaller oligomers and even dimers also exist in model membranes ^19,31–34^. The intrinsic fluorescence of nystatin and natamycin is based on four conjugated double bonds, and we have shown recently, that the interaction of nystatin with cell membranes can be followed by ultraviolet (UV) sensitive microscopy ^35^. We found that nystatin binds preferentially to yeast cells containing ergosterol compared to those loaded with cholesterol ^35^. We observed highly fluorescent nystatin aggregates at the surface of mammalian cells and yeast cells loaded with cholesterol but much less so for yeast cells containing ergosterol ^35^. Surface-attached nystatin aggregates were inefficient in extracting the fluorescent cholesterol analogue TopFluor-cholesterol from mammalian cells, arguing against the extracellular sponge model for the action of this polyene ^12,29,35^. Whether natamycin forms aggregates in yeast cells as part of its antifungal activity is not known.

Here, we make use of the intrinsic fluorescence of natamycin to visualize its interaction with yeast cells. By combining fluorescence and X-ray microscopy we show that natamycin forms aggregates in the yeast PM in a sterol-dependent manner and causes leakage to propidium iodide (PI) in about 2.5h. Aggregation of natamycin in the yeast PM is not the result of a clustered sterol distribution, since intrinsically fluorescent, close analogs of both, ergosterol and cholesterol, have a homogeneous lateral distribution in the yeast PM. Depletion of sphingolipids using the synthesis inhibitor myriocin increased natamycin binding, likely by redistributing ergosterol from the inner to the outer PM leaflet. Thus, the ergosterol-dependent structural organization of the yeast PM causes the preferred binding of natamycin and can explain the selective antifungal activity of this polyene. Our results thereby set the stage for future studies directed towards mechanistic exploration of natamycin’s mode of action against pathogenic fungal species.

## Results

### Quantitative fluorescence imaging of binding and uptake of natamycin

Due to its four conjugated double bonds, natamycin has an intrinsic fluorescence with an excitation maximum at 305 nm and a broad emission spectrum with a maximum at 404 nm ^17^. Employing UV-sensitive fluorescence microscopy with excitation and emission filters matching the spectra of natamycin, it is possible to visualize binding of the polyene to membranes (see Materials and methods). We have used this approach recently to show that natamycin binds to the liquid-ordered phase in giant unilamellar vesicles (GUVs), particularly in those GUVs containing ergosterol ^18^. Using the same approach in living yeast cells, we find that natamycin binds initially to the PM, containing the eisosome marker YPet-tagged Sur7 (Sur7-YPet) (Fig. 1A) ^2^. To determine, how binding of natamycin to yeast cells is related to the ability of the polyene to kill yeast, we have implemented multi-color time-lapse microscopy, in which yeast expressing Sur7 was incubated with natamycin in the presence of PI, a membrane impermeable DNA stain. PI emits red fluorescence and only label cells, when intactness of the PM is compromised, i.e., in dead or dying cells ^36^. By three-color time-lapse microscopy, we find that PI gets access to the cell interior resulting in red intracellular fluorescence after 140-170 min incubation with natamycin (Fig. 1A). Natamycin stays at the PM for up to 2h, before the polyene slowly gets internalized (Fig. 1B). This was inferred from a decrease in peripheral and an increase in intracellular staining surrounding the vacuole (Fig. 1B). After 6h, natamycin and Sur7-YPet have moved inward (not shown), and after 16h incubation, both are found inside the cells, where they co-localize with the vacuole marker FM4-64 (Fig. 1C). Together, these results show that the PM is the target site for the polyene, and once the integrity of the cell membrane is compromised, natamycin enters the cell and stains internal compartments, such as the vacuole. Thus, even though natamycin is not known to form pores for ions in membranes ^13,16,28^, it changes the membrane integrity and facilitates passage of small molecules, like PI. This finding is in accordance with our recent observation, that access of the aqueous quencher dithionite to the fluorescent phospholipid analogue NBD-PC is increased in the presence of natamycin in model membranes ^18^.

**Figure 1.**
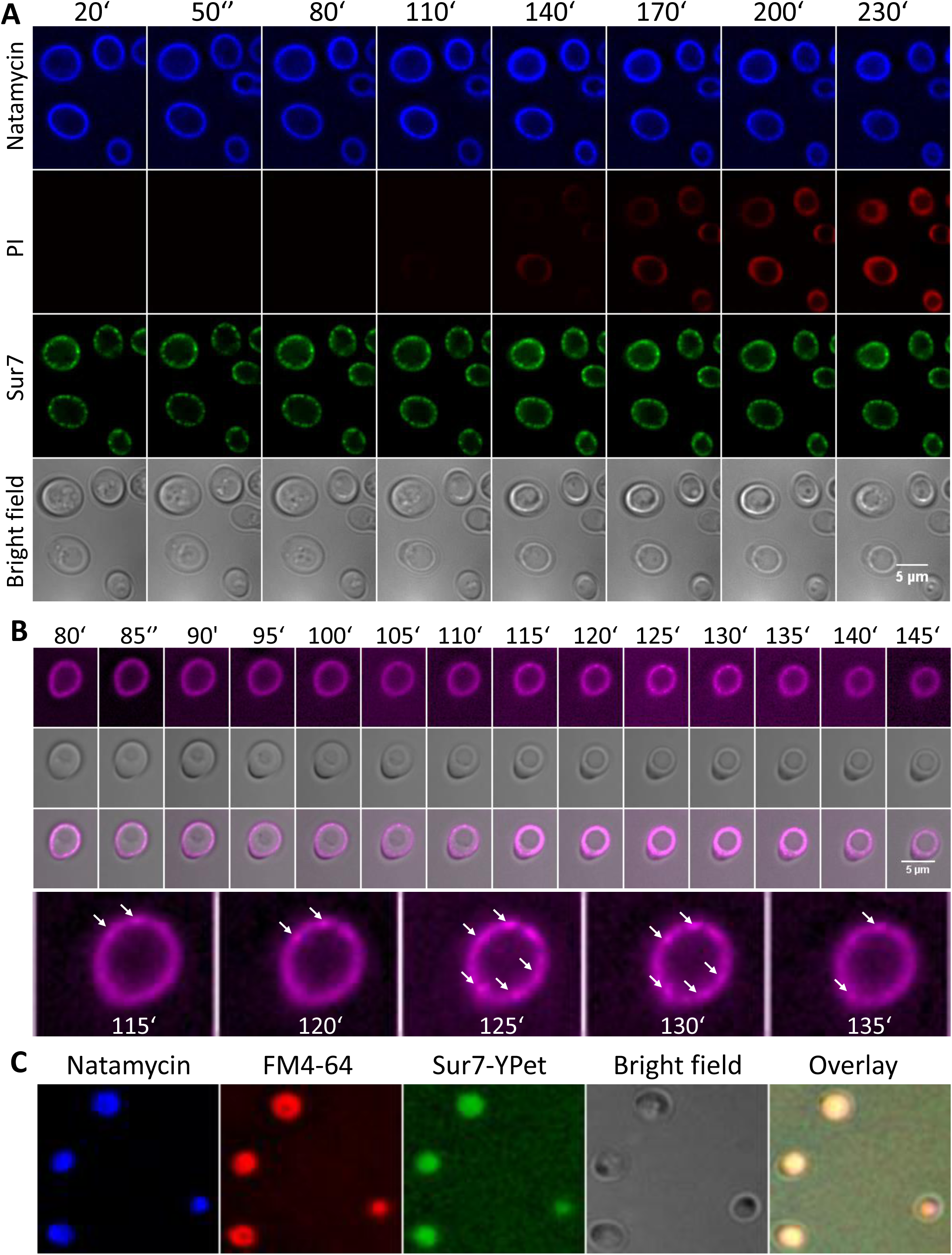
Kinetics of binding and membrane activity of natamycin by time-lapse microscopy. A, Ypet-sur7 yeast cells were incubated with 50 µg/ml natamycin, washed briefly and imaged on a UV-sensitive widefield microscope every 30 min. The dye propidium iodide (PI) was provided in a PBS solution throughout the incubation period on the microscopy dish. Scale bar, 5 µm. B, yeast cells were incubated with natamycin for the indicated time, washed and imaged by UV-sensitive time-lapse microscopy. Spots of increased natamycin intensity were detected in the PM between 115 and 135 min of incubation (arrows). Cell entry was observed after ca. 140 min, when the diameter of the ring-like staining became smaller than the cell diameter. Scale bar, 5 µm. C, cells expressing YPet-Sur7 (green) were incubated with natamycin (blue) and with the vacuole marker FM4-64 (red) for 16h, washed and imaged. The color overlay shows co-localization in the vacuole in yellow.

### Natamycin forms clusters in the yeast PM as visualized by fluorescence and X-ray microscopy

The results shown above give us a time window of about 2.5h to study the impact of natamycin on the yeast PM before cellular integrity is compromised. By live-cell imaging, we observed formation of natamycin patches in the PM right before uptake, i.e., after 115 to 130 min of incubation but long after its initial binding to the yeast PM (Fig. 1B, arrows). This could indicate preferred binding to some membrane areas or formation of natamycin aggregates in the PM. Supporting an aggregation model, we found previous evidence for the formation of natamycin oligomers in aqueous solution and model lipid membranes based on alterations in the fluorescence spectrum and on molecular dynamics (MD) simulations ^17^. Here, we observe that after 2h of incubation of yeast with natamycin, lateral intensity peaks were pronounced resulting in heterogeneous PM staining (Fig. 2). This was assessed by extracting line profiles along the membrane, from which the coefficient of variation (CV) i.e., the signal’s standard deviation divided by the mean is calculated (see Fig. S1A and Materials and methods for the procedure). We verified in simulations that the CV is a good measure of heterogeneous line profiles (see Materials and Methods and Fig. S1B-G). In particular, the CV depends linearly on the amplitude of intensity variations which is directly related to the size of clusters (Fig. S1F and G). Since we pre-processed all images by 2D Richardson-Lucy deconvolution before analysis (see Materials and methods), we also assessed the impact of this method on our quantification protocol. We found that deconvolution causes a non-linear increase in the CV values of line profiles of natamycin-stained cells as a function of iteration (Fig. S2). This is a direct consequence of the increased signal-to-noise ratio which results in larger amplitude variations upon deconvolution. The CV measure becomes independent of deconvolution iteration, once the algorithm converges, which is after about 30 iterations. Thus, we deconvolved all images by the same procedure with 30 iterations of the Richardson-Lucy algorithm. The large CV values we obtained for natamycin-treated cells indicate that polyene aggregates form at or in the yeast PM. The extent of clustering of natamycin varied significantly between yeast cells, suggesting that aggregation of the polyene in the PM is a stochastic event (Fig. 2B-F).

**Figure 2.**
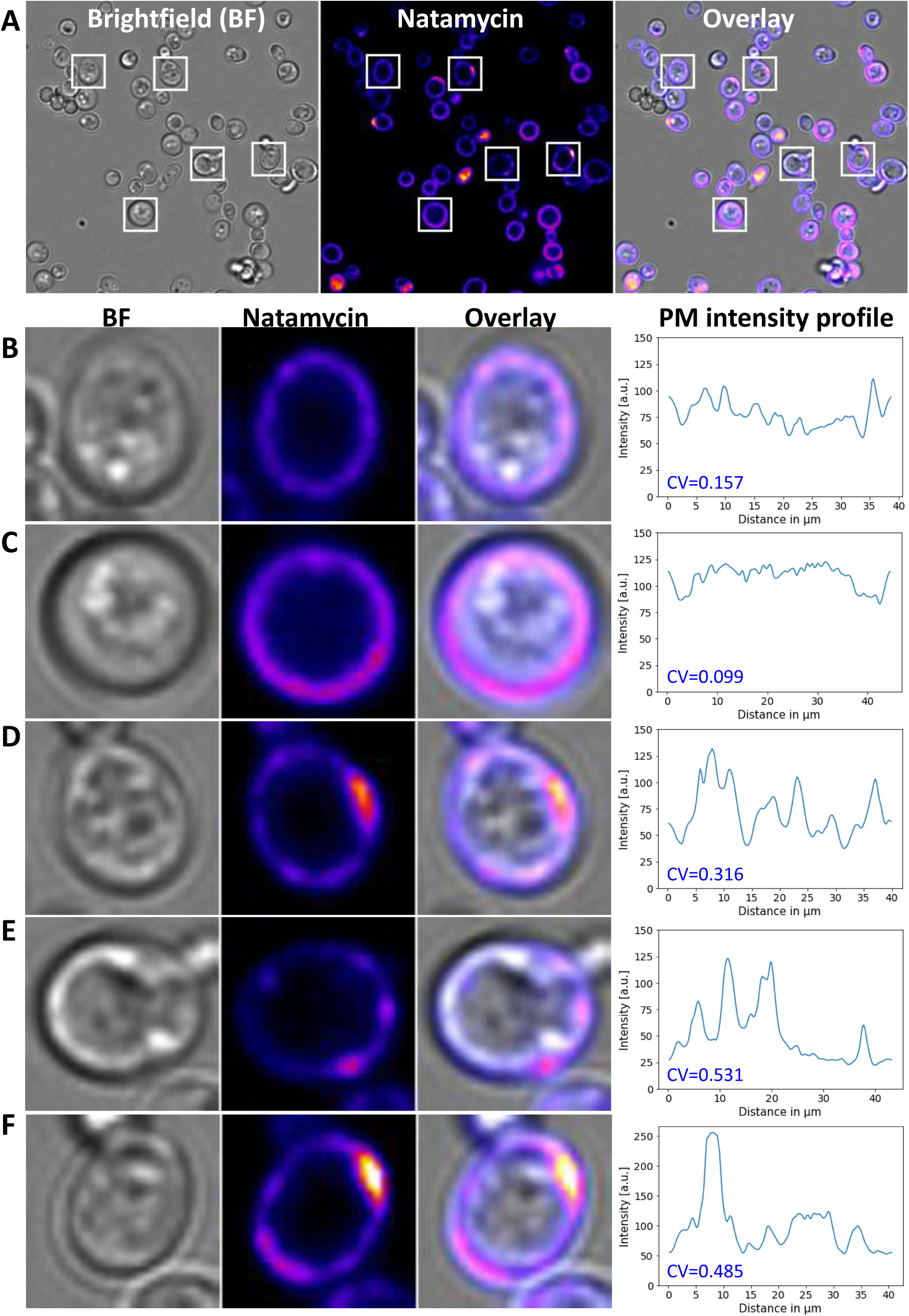
Natamycin forms clusters in the PM prior to entry into cells. A, Cells were incubated with natamycin (final concentration of 50 µg/ml) for 2h prior to imaging which leads to natamycin clustering in the PM for cells. B-F, images of cells highlighted in A. B-F right panels, extracted line profiles of cells used for calculating the coefficient of variation (CV). See main text for details.

While we did not find evidence for significant natamycin aggregation in the cell medium, UV-sensitive fluorescence microscopy lacks the resolution to definitely test an aggregation model and also to rule out that the observed aggregates formed at the yeast cell surface, i.e., attached to the outer side of the cell wall. To study the impact of natamycin on yeast cells at the ultrastructural level, we used soft X-ray tomography (SXT), a method that that uses the absorption of X-rays in the so-called water window by carbon-rich material as contrast principle ^38^. SXT allows for imaging entire yeast cells in their near-native state upon cryo-fixation and without the need for cell fixation ^39–41^. Using a synchrotron as source of X-rays in the appropriate energy range of about 510 eV and suitable zone plates to focus the incident X-rays to the sample, SXT can obtain 30-50 nm isotropic resolution with typical acquisition times of about 30 min per tomogram ^42,43^. Due to their high abundance of carbon, natamycin aggregates forming in solution above the critical aggregation concentration of the polyene ^17^ can be easily visualized by SXT (Fig. S3). In cell samples incubated with natamycin, such aggregates of heterogeneous shape and diameter were occasionally observed embedded in the ice but at distant locations from cells and not attached to the cell wall (not shown). This excludes, that the natamycin clustering, we observe in fluorescence microscopy stems from surface-attached aggregates. Treating cells with natamycin caused deformations and occasional blebbing of the PM and cell wall, as clearly seen in SXT reconstructions (Fig. 3A). In addition, we observed dark structures resembling polyene aggregates underneath the cell wall of natamycin-treated cells in the reconstructed X-ray tomograms, both underneath the cell wall and in association with the vacuole (Fig. 3B, C, white arrows). Such structures were in the size range of 60 to 170 nm and were not observed in non-treated cells (Fig. S3). To confirm the identity of such structures as natamycin aggregates, we also carried out X-ray microscopy of pre-formed aggregates formed in aqueous solution (Fig. S3F). This experiment showed that aggregates of natamycin strongly absorb X-rays, and it revealed that such aggregates are of heterogeneous size and shape, with the smallest structures being observed in the size range as found in cells.

**Figure 3.**
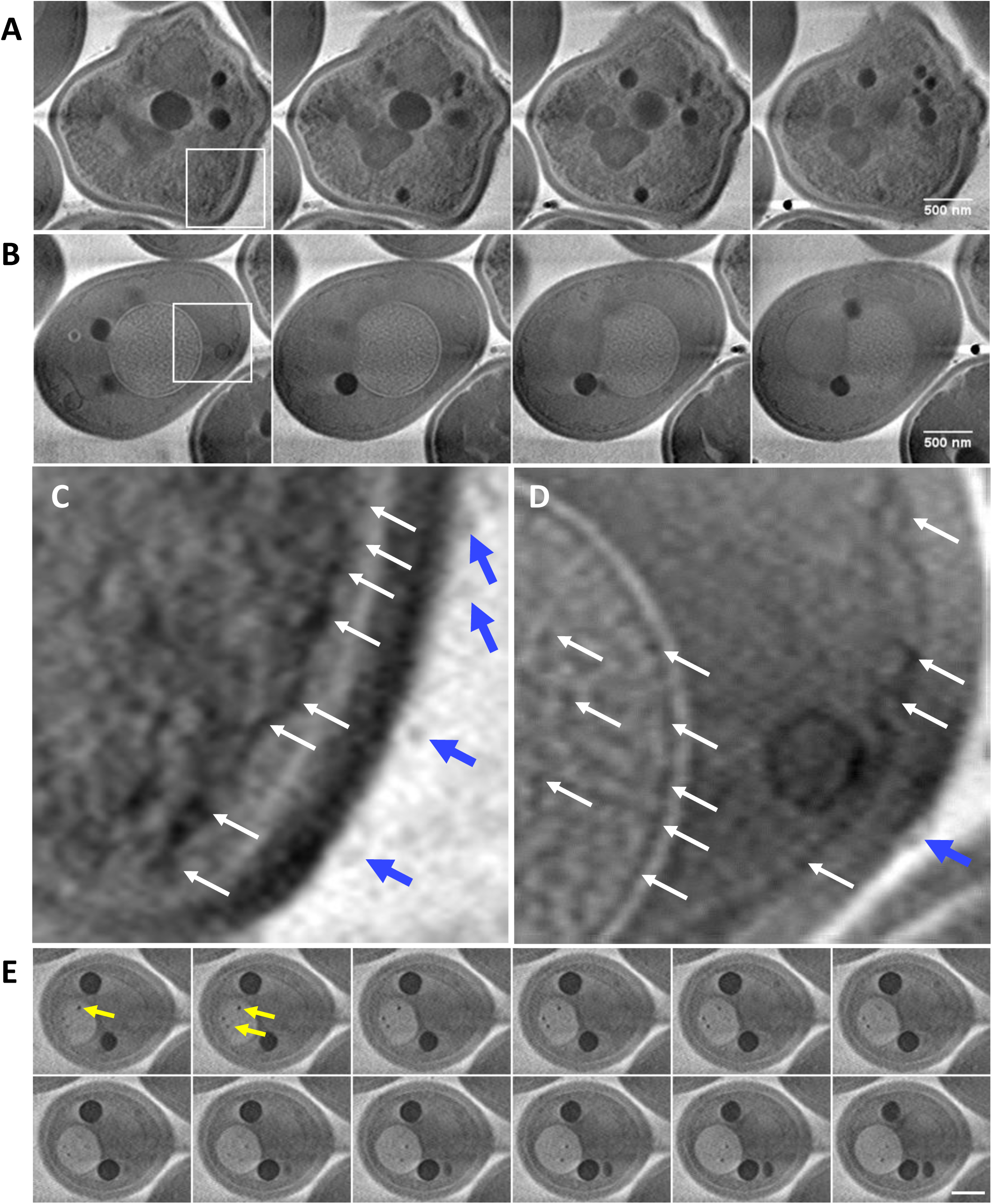
Soft X-ray tomography of natamycin-treated yeast cells. SXT was carried out on plunge-frozen yeast cells after incubation with PBS buffer or with PBS containing 50 µg/ml natamycin for 2h. Tomograms were aligned and reconstructed as described in Materials and methods. A, B, montage of selected frames from two 3D reconstructions, showing deformed cell shape and membrane detachment upon natamycin treatment. C, D, zoomed version of panels A and B, respectively, showing dark aggregates underneath the cell wall (C, white arrows) and PM detachment/invaginations (D, white arrows). Outside of the cells, extracellular vesicles were found, which locate closely to the cell wall (C, D, blue arrows). E, montage of another natamycin-treated cell with polyene aggregates inside of the vacuole (see yellow arrows in the first two frames pointing to dark spots inside the vacuole). Bar, 0.5 µm.

X-ray microscopy also revealed the ultrastructural changes of yeast upon natamycin treatment, including membrane detachments and formation of extracellular vesicles (Fig. 3C, D, blue arrows). The lipidic nature of such structures could be confirmed by staining with Nile Red, a membrane inserting hydrophobic dye, using confocal microscopy (not shown). Formation of such extracellular vesicles could represent a first line of defense to get rid of the drug molecules, as has been shown in *C. neoformans* upon azole treatment ^64^. Distinct aggregates with a size of 350 to 500 nm were occasionally also observed inside the vacuole by SXT (Fig. 3D and E). This confirms our fluorescence imaging results (compare Fig. 1C and Fig. 3E). We conclude that small natamycin aggregates form initially in association with the yeast PM, either as membrane-attached structures or upon intercalation into the bilayer. This causes extensive membrane deformation, detachment from the cell wall and release of extracellular vesicles. After prolonged exposure of cells, natamycin also accumulates in intracellular organelles, such as the vacuole.

### Natamycin clusters in the yeast PM with partial preference for MCC/eisosomes

To determine, whether natamycin aggregates preferentially in eisosomes, we used our direct imaging approach to quantify the lateral distribution of bound natamycin compared to known protein markers in the yeast PM. Sur7-YPet is an established marker for eisosomes/MCC ^7^, and we measured the intensity profile of natamycin and Sur7-YPet in the PM of aligned images of double-labeled cells to determine eventual co-patching of natamycin with this eisosome marker, (Fig. 4A and B). There is some co-localization between the polyene and Sur7-YPet as inferred from the Pearson correlation coefficient, which is *r*=0.54 for the profile shown in Fig. 4A and B. We validated this correlation analysis by carrying out a Bayesian estimate of the correlation between both intensity profiles (Fig. 4B-I). Assuming a bivariate normal distribution for both intensity profiles, the Bayesian analysis provides the complete probability density function (PDF) of the estimated correlation coefficient (i.e., the posterior distribution, Fig. 4C). For that, Monte Carlo sampling of the posterior distribution is carried out using the NUTS sampler in PyMC3, a python library for Bayesian inference of experimental data ^44^. This analysis reveals that 94% of all correlation coefficients for this data set lie between *r*=0.34 and 0.69 with a mean value of *r*=0.52. Thus, the simple calculation of the Pearson coefficient and the mean of the Bayesian equivalent are comparable, and the PDF estimated by the Bayesian analysis is rather narrow. The Bayesian analysis not only provides the PDF for the correlation coefficient, but also the distributions of the mean intensities and the width of such distributions and thereby allows for simulating intensity profiles based on the experimental measures for both, natamycin and Sur7-YPet (Fig. 4D-I). From that, one sees that for the given example, the mean intensity of natamycin is higher than for Sur7-Ypet but the intensity variation is larger for the eisosome marker (compare Fig. 4E and I and see Fig. S4 for another example). Since the Bayesian method is computationally expensive, it cannot be employed for the entire experimental data set of double-labeled cells. Therefore, for a comparative analysis of many cells, we only used the simple Pearson coefficient to assess the correlation between membrane staining of natamycin and Sur7-YPet. The distribution of Pearson coefficients measured between intensity profiles of natamycin and Sur7-YPet for more than 100 cells is multimodal and spans a large range of correlation values (Fig. 4J, upper bar chart). We employed a Gaussian mixture model to dissect this multimodal distribution, identifying three populations of Pearson values (Fig. 4J). The first population resembles cells with high correlation, the second with low, and the third with no correlation between both intensity patterns. Thus, natamycin tends to co-patch with Sur7-YPet in about 1/3 of the cell population, while it shows little co-patching in about 50% of the cells and no correlation or even slight anti-correlation in the remaining cells. The underlying causes of this cell-to-cell heterogeneity are not clear at the moment, but we surmise, that the exact aggregation pattern of natamycin depends on the stochastic event of aggregate initiation at or in the PM. Once small clusters are formed, larger aggregates might form by addition of further natamycin monomers, and this process could happen inside or outside of MCC/eisosomes depending on the initial oligomerization event. While this model is difficult to prove given the current resolution of the used microscope systems, we do find extensive co-clustering of natamycin inside as well as outside of MCC/eisosomes even within the same cell (Fig. S4). Also, MCC/eisosomes are membrane invaginations and thus there is more membrane surface in 2D views of MCC/eisosomes than for the rest of the membrane (and thus there is more room for natamycin partitioning). Accordingly, if initial association of natamycin happens to take place in MCC/eisosomes, further aggregation in these compartments is likely. Supporting a model of stochastic association of natamycin with membrane domains, we found , also partial co-localization with the MCP marker Pma1-YPet (Fig. S5). In conclusion, our results demonstrate that aggregation of natamycin does not strictly depend on the location of MCC/ eisosomes or other protein domains in the yeast PM.

**Figure 4.**
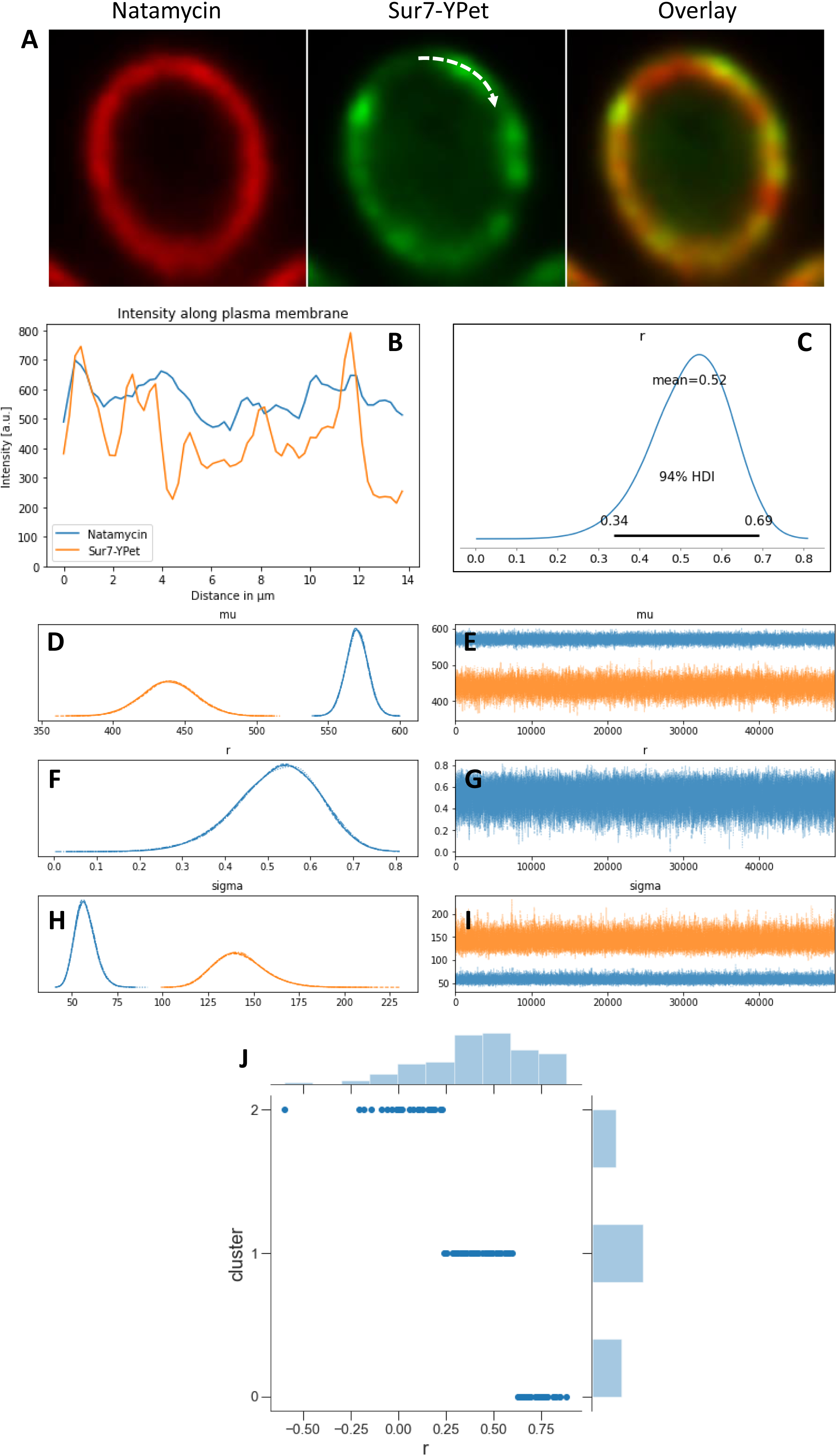
Natamycin clusters form inside and outside of eisosomes in the yeast PM. Yeast cells expressing Sur7-YPet, a green-fluorescent eisosome marker, were incubated with natamycin for 2h, washed and imaged. A, natamycin pseudo-colored in red, Sur7-YPet (green) and color overlay; the dashed arrow in the middle panel indicates the start of the extracted line profile. B, line profiles for natamycin (blue line) and for Sur7-YPet (orange line) for the cell shown in A. C, probability density function (PDF) of Pearson coefficient, *r*, estimated by a Bayesian analysis with mean (r=0.52) and the highest density interval (HDI). D, Bayesian estimate of the PDF of mean intensity values for each marker, i.e., posterior for Sur7-YPet (orange line) and natamycin (blue line) compared to the respective priors (dashed lines). E, corresponding MCMC runs for both intensities. F, comparison of posterior (straight line) with prior (dashed line) of the Pearson coefficient. G, corresponding MCMC run for the Pearson estimates. H, prior (dashed lines) and posterior distribution (straight lines) for the standard deviation of the intensities of Sur7-YPet (orange lines) and natamycin (blue lines). I, corresponding MCMC runs as function of iteration. J, Pearson coefficients were calculated for the entire population of analyzed cells (n=110) and deconvolved into three populations using a Gaussian Mixture model. See main text for further information.

### Binding and clustering of natamycin in the yeast PM is sterol-dependent

To determine, whether binding and clustering of natamycin depends on sterols, we made use of a yeast strain, in which ergosterol synthesis is blocked, by a mutation in heme synthesis (*hem1*Δ cells) ^45^. In such cells, sterol import via the ABC transporters Aus1 and Pdr11 is upregulated to ensure sterol supply from the culture medium ^46–48^. This system allows us to load cells with a given sterol under normal aerobic growth conditions and thereby to assess the sterol dependence of polyene binding. Binding of natamycin was strictly dependent on sterols in the PM, as in *hem1*Δ cells incubated in the absence of sterols, binding of natamycin was abolished (Fig. S5). Similarly, in a triple yeast mutant strain lacking the sterol importers Aus1 and Pdr11 (*hem1*Δ/*aus1*Δ/*pdr11*Δ cells) binding of natamycin to the PM was strongly diminished (Fig. S6). These results show that sterol import into the PM by the ABC transporters Aus1/Pdr11 is instrumental for natamycin binding under conditions of blocked ergosterol synthesis. Passive association of sterols with the cell wall is not sufficient for natamycin’s interaction with the yeast PM. Both, the yeast sterol ergosterol and mammalian cholesterol can mediate binding of natamycin to yeast cells, but binding of the polyene was much stronger, when cells were loaded with ergosterol compared to cholesterol. This was first concluded from measurements of cell-associated natamycin intensity (Fig. S6). In independent experiments, we measured line profiles of natamycin-stained cells along the PM, as described above, and found that the mean of the intensities of ergosterol-loaded cells after background subtraction were 354.1± 98.9, whereas for the cholesterol-loaded cells we found a value of 268.7± 47.2, i.e., ca. 76% of the natamycin intensity in ergosterol-containing cells. Thus, natamycin binds preferentially to yeast cells containing the native yeast sterol ergosterol.

From the measured intensity profiles, we also determined the tendency of natamycin to cluster in the PM of yeast loaded with either ergosterol or cholesterol. We found that the extent of natamycin aggregation in the PM is larger in cells loaded with ergosterol than with cholesterol (Fig. 5). To quantify this observation, an automated analysis of cluster sizes was carried out based on the extracted line profiles using a peak finding algorithm (Fig. 5E-H). This analysis revealed that the clusters were of more distinct size in cells loaded with ergosterol compared to those loaded with cholesterol, in which the aggregate size of natamycin scattered widely (Fig. 5E-H). The average cluster size was 516 nm ± 29.9 nm (n=394 cells from 3 experiments) in ergosterol-loaded cells compared to 464 nm ± 37.4 nm (n=305 cells from 3 experiments) in cholesterol-loaded cells (Fig. 5G). Larger cluster sizes are also reflected in higher CV values for ergosterol-compared to cholesterol-containing membranes (Fig. 5H). Since we also find comparable uptake of intrinsically fluorescent analogues of both sterols (see below), the observed differences in natamycin binding are not due to differing loading efficiencies with the sterols. We thus conclude that aggregation of natamycin is less pronounced in the PM of cholesterol-containing yeast compared to yeast containing ergosterol. It is likely that coalescence of smaller natamycin oligomers into larger aggregates takes place primarily in ergosterol-containing membranes, while natamycin clusters remain small in cholesterol-containing yeast cells. Together, these results show that sterols are essential for natamycin’s interaction with yeast cells. Ergosterol is clearly more efficient in recruiting natamycin to cells and in mediating aggregation of natamycin in or at the PM compared to cholesterol. These results provide a mechanistic basis for the selective antifungal activity of natamycin.

**Figure 5.**
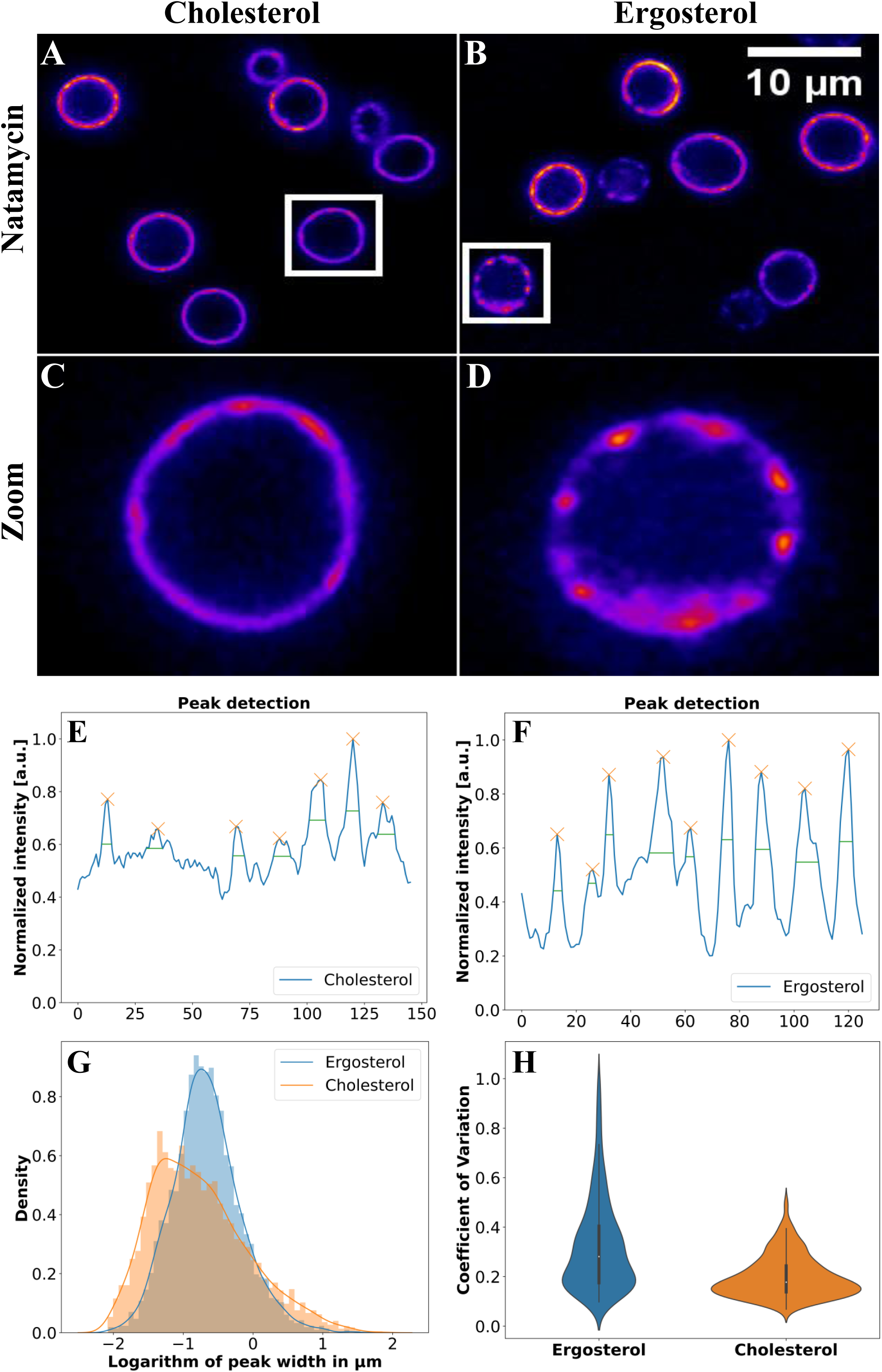
Natamycin clusters in the PM of yeast in a sterol-specific manner. *hem1*Δ cells were grown in media supplemented with 5 μg/ml cholesterol (A,C) or ergosterol (B, D) for 22h and treated with natamycin (final concentration of 50 µg/ml) for 2h. Scale bar, 10 µm. E-F, peak detection obtained from the line profile of yeast labelled with cholesterol (C) or ergosterol (D) normalized to the maximum value. Peaks are highlighted with orange crosses and the selected width of the peak in green. G, density plot of the logarithm of peak width in µm for cells with cholesterol (orange) or ergosterol (blue). H, violin plot of the coefficient of variation (CV) for cells loaded with cholesterol (n=305) or ergosterol (n=394).

### Inhibition of sphingolipid synthesis acts synergistically with natamycin

The yeast PM contains specific sphingolipids with long saturated chains that are not found in mammalian cells, such as inositol-phosphoryl ceramides, which means that sphingolipid synthesis represents an important drug target in pathogenic yeast species ^49^. Myriocin is an inhibitor of the first step in sphingolipid synthesis, the condensation of serine and palmitoyl-CoA catalyzed by serine-palmitoyl transferase. Thus, treating cells with myriocin abolishes the production of sphingolipids entirely. Strikingly, when incubating yeast with myriocin before treatment with natamycin, the number of PI-stained cells was strongly increased (Fig. 6A-D). At the same time, cell-associated fluorescence of the polyene was increased by about 30% compared to cells incubated in the absence of myriocin (Fig. 6E-G). This suggests? that more ergosterol may be accessible for recruiting natamycin to the PM in the absence of sphingolipids, likely because ergosterol redistributes from the inner to the outer PM leaflet. This conclusion is supported by our previous study, in which we used side-specific quenchers of the intrinsically fluorescent ergosterol analogue DHE to show that most sterol is in the inner PM leaflet of yeast cells with only about 22% of PM sterol residing in the outer lamella of the cell membrane ^10^. This fraction increased to almost 45% upon treating cells with myriocin, an inhibitor of sphingolipid synthesis ^10^. Thus, binding of natamycin to yeast cells depends on the accessibility of ergosterol to the polyene, which likely resembles the ergosterol pool in the outer PM leaflet. This sterol pool is increased upon myriocin treatment without altering DHE uptake into *hem1*Δ cells, as we showed previously ^10^. We conclude that inhibitors of sphingolipid synthesis can act synergistically with natamycin in killing yeast, likely by altering the transverse sterol distribution in the PM thereby enhancing natamycin binding to cells.

**Figure 6.**
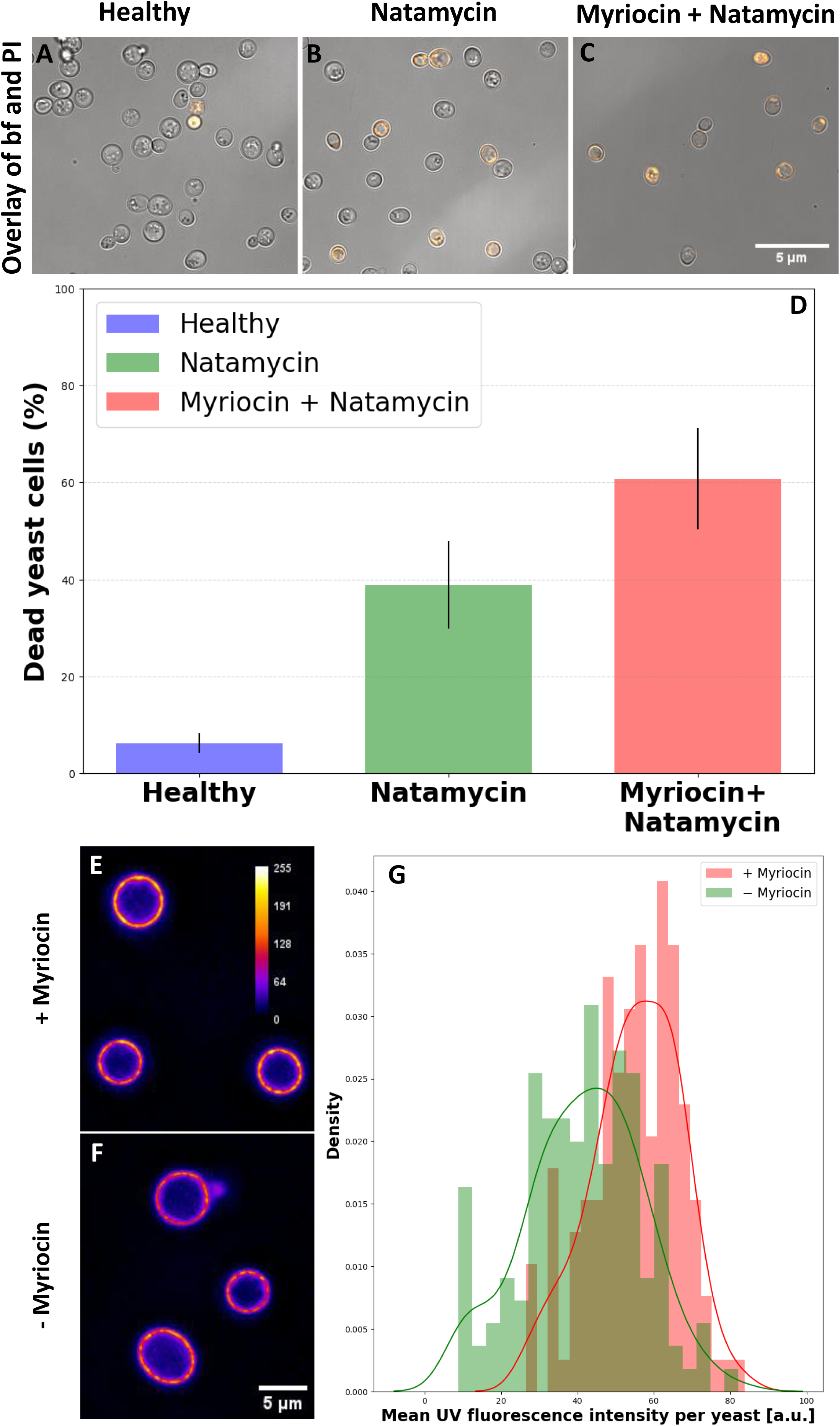
Inhibition of sphingolipid synthesis can act synergistically with natamycin in killing yeast. *hem1*Δ cells were grown in media supplemented with *δ*-aminolevulinic acid (dALA) to ensure heme synthesis and thereby ergosterol biosynthesis. To inhibit sphingolipid synthesis cells were treated with myriocin and natamycin for 2h. Cells were stained with propidium iodide (PI) for 5 min before imaging. A-C, overlay of brightfield and PI channel for cells without any treatment (A), treatment with natamycin (B), and double treatment with myriocin and natamycin (C). Scale bar, 5 µm. D, for each condition (A-C) the percentage of dead cells was quantified as being PI-positive upon cell segmentation using Cellpose (see Material and methods). E, F, example image of cells treated with myriocin and natamycin (E) compared to cells treated with natamycin only (F). G, density plot displaying the mean UV fluorescence intensity per yeast (a.u.) for cells treated with myriocin and natamycin (red; n=137) and only natamycin (green; n=150).

### Analogues of ergosterol and cholesterol have a homogeneous lateral distribution in the PM

To rule out that the higher aggregation propensity of natamycin in the PM of yeast loaded with ergosterol compared to that in cholesterol-loaded cells is a consequence of more efficient ergosterol uptake by Aus1/Pdr11, we loaded cells with the intrinsically fluorescent analogues of either ergosterol (DHE) or cholesterol, (CTL). DHE and CTL differ from ergosterol and cholesterol only by one or two double bonds in the steroid ring system (Fig. 7), respectively. They differ from each other only in their alkyl side chain, which is that of ergosterol for DHE and that of cholesterol for CTL. We found that cell-associated intensities of DHE and CTL were comparable, showing that cells take up both sterols to a similar extent (Fig. 7A-D). Quantification of the intensity of both sterols in the PM from extracted line profiles even found slightly higher staining with CTL compared to DHE (i.e., 279.0 ± 39.2 for DHE and 311.9 ± 40.8 for CTL). Insertion of both sterols into the PM required Aus1/Pdr11, as it was blocked in cells lacking these transporters (not shown but see ^47^). These results show that minimally modified analogs of ergosterol and cholesterol are equally efficient in yeast via the Aus1/Pdr11 transport system when ergosterol synthesis is blocked. Accordingly, the higher membrane association we found for natamycin in ergosterol-loaded cells is due to the preferred interaction of the polyene with the yeast sterol compared to mammalian cholesterol.

**Figure 7.**
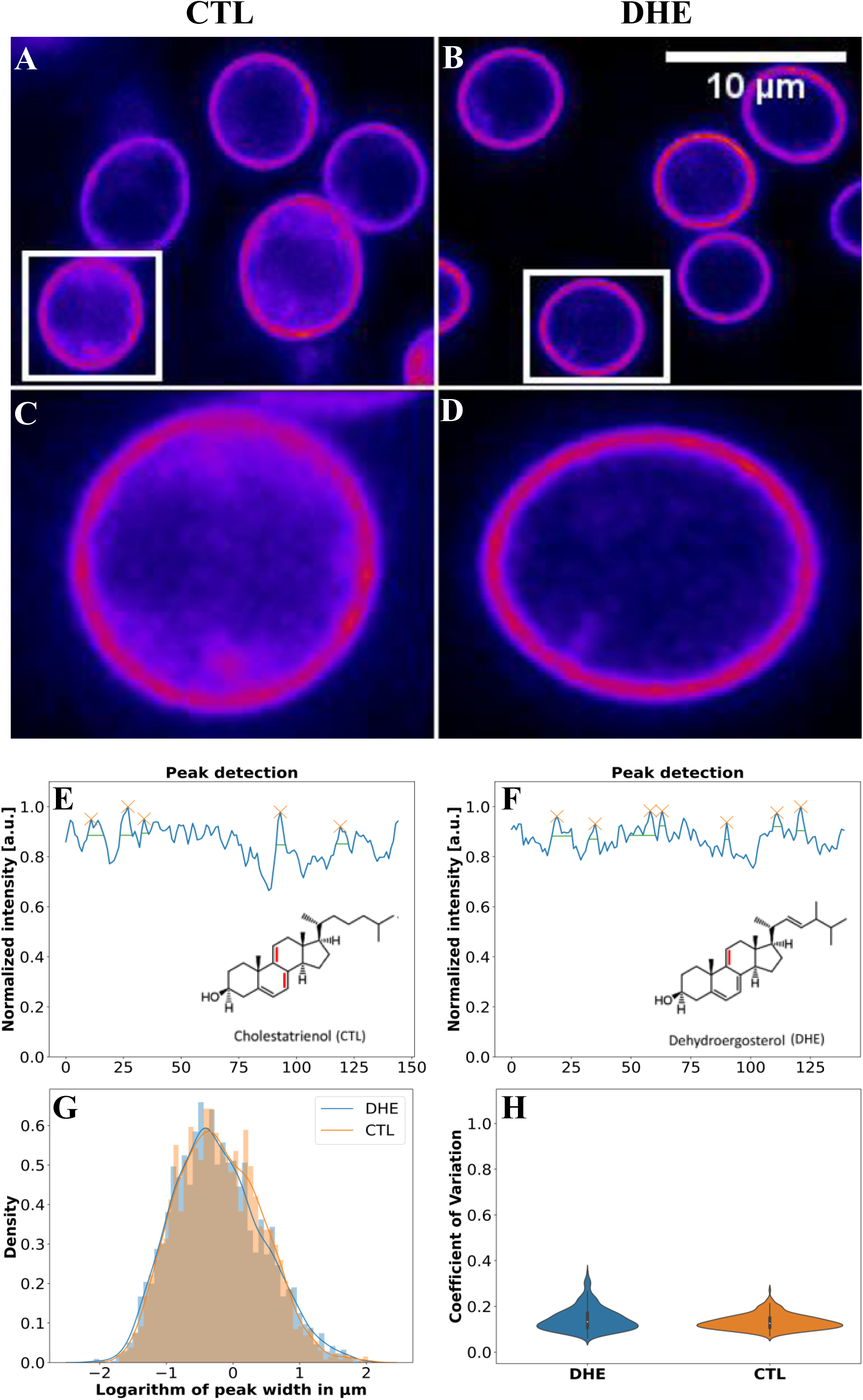
Analogues of ergosterol (DHE) and cholesterol (CTL) have a homogeneous lateral distribution in the PM. *hem1*Δ cells were grown in media supplemented with 5 μg/ml CTL (A,C) or DHE (B, D) for 22h. Scale bar, 10 µm. DHE and CTL differ from ergosterol and cholesterol only by one or two double bonds in the steroid ring system which provides the sterols with fluorescence properties emitting light in the UV region of the spectrum highlighted in red in E and F, respectively. E-F, peak detection obtained from the line profile of yeast labelled with CTL (C) or DHE (D) normalized to the maximum value. Peaks are highlighted with orange crosses and the selected width of the peak in green. G, density plot of the logarithm of peak width in µm for cells incubated with CTL (orange) or DHE (blue). H, violin plot of the coefficient of variation (CV) for cells loaded with CTL (n=300) or DHE (n=551).

The heterogeneous distribution of natamycin raises the question of how sterols, which are essential for natamycin binding to yeast cells, are laterally distributed in the PM. Work in the Tanner lab used filipin to show a preferred staining of eisosomes and concluded that these are ergosterol-rich compartments ^2,50^. Others could not confirm these results with filipin ^51^, and more recent studies relying on biochemical analysis of isolated membrane patches found a rather even distribution of ergosterol between different domains of the yeast PM ^8^. By applying our analysis to yeast incubated with the fluorescent sterols, we found a largely homogeneous distribution of both, DHE and CTL in the yeast PM (Fig. 7C and D). This is reflected in small intensity fluctuations with much lower CV values for both sterols compared to natamycin (compare Fig. 6E, F with Fig. 7E, F). We conclude that these intensity variations are largely due to topographic inhomogeneities of the yeast PM, similar to those previously shown for mammalian cells ^52,53^. Together, our results demonstrate that close analogues of ergosterol and cholesterol have a rather homogeneous lateral distribution in the yeast PM. Consequently, we can conclude that natamycin does not bind to pre-existing sterol domains but rather forms aggregates once it is associated with the cell membrane.

## Discussion

In the present study, we established a method to visualize the interaction of natamycin with yeast cells based on the polyenes’ intrinsic fluorescence. This novel approach allows us to directly observe the interaction of natamycin with yeast without the need for labeling, which can compromise the molecular properties of polyene drugs ^54,55^. We demonstrate that natamycin rapidly binds to yeast cells and interacts with the PM in a sterol-dependent manner (Fig. 2 and 5). Using an ergosterol-deficient yeast strain with defective sterol synthesis, we show that natamycin binds preferentially to yeast cells loaded with ergosterol compared to yeast loaded with cholesterol, and we show that this is not due to preferred ergosterol uptake (Fig. 5 and S6). Using NMR spectroscopy of deuterated sterols and MD simulations, we demonstrated recently that natamycin immobilizes both sterols in model membranes to a similar extent ^18^. At the same time, we observed that natamycin interacts preferentially with ergosterol-containing model membranes interfering with lipid phase properties and transport activity of reconstituted amino acid transporters ^17,18^. Based on these findings, we conclude that the preferred interaction of natamycin with ergosterol-containing yeast must be due to specific properties of the PM mediated by the yeast sterol, which cannot be replaced completely by mammalian cholesterol. This conclusion is in accordance with our previous spectroscopic studies of natamycin in model membranes and also in agreement with studies by others on membrane interaction of the related nystatin ^56–58^. On the other hand, for nystatin, we found an even more pronounced ergosterol dependence of binding to yeast cells using our label-free imaging approach ^35^. Both, nystatin and AmpB form sterol-dependent ion channels in membranes, while natamycin does not, and for AmpB ergosterol-specific interactions in lipid bilayers have been demonstrated using NMR spectroscopy ^13,59,60^. Such divergent findings indicate fundamental differences in the mechanism by which various polyene molecules attack lipid membranes. That yeast imports sterols from the medium via Aus1/Pdr11 when the synthesis of ergosterol is blocked can explain why some pathogenic yeast strains become resistant to azoles, such as fluconazole, known to inhibit ergosterol biosynthesis ^61^. Our results suggest that treatment of such cells with natamycin or other polyenes, like nystatin, could overcome the drug-resistant phenotype since binding of the polyenes to the PM is reduced but not prevented in the presence of cholesterol imported from the host cells.

By combining label-free fluorescence with X-ray microscopy we show that natamycin forms small aggregates in or at the PM, and this process is enhanced in cells containing ergosterol compared to those containing cholesterol. These findings demonstrate that membrane-associated aggregation of natamycin is a key aspect of its antifungal activity. Whether such natamycin aggregates form inside of the PM bilayer or on top of the cell membrane, as recently proposed in the sterol sponge mechanism for the mode of action of glycosylated polyene macrolides remains to be shown ^12,29^. Given that we find aggregates also inside the cells including the lumen of the vacuole after prolonged incubation, indicates that formation of polyene sponges is not restricted to the cell surface. Using time-lapse microscopy, we show that lateral clustering of natamycin in the PM happens after its initial binding to the cells, making cell association of pre-assembled natamycin aggregates unlikely (Fig. 1). However, our data does not exclude that some of the smaller aggregates can pass through the cell wall and associate with the yeast PM independent of the membrane association of natamycin monomers. Supporting a model of polyene action from inside the PM, label-free imaging of AmpB aggregates revealed their residence in yeast and mammalian cell membranes and polarization-sensitive Raman imaging found an orientation of this polyene parallel to the bilayer normal in yeast cells ^32,65,66^.

Our time-lapse data also shows that aggregation of natamycin in the PM precedes permeation of the small molecule PI into cells (Fig. 1). In model membranes, we observed recently that natamycin increases permeation of dithionite across the bilayer, supporting the conclusion that it can compromise membrane integrity ^18^. Given the timeline, we hypothesize that aggregate formation is a precondition for membrane leakiness by natamycin. Natamycin does not form aqueous pores in membranes ^13,16^, and we confirm that by challenging cells with hyperosmotic shocks, where we found that natamycin-treated cells are in fact slightly protected against osmotic stress (Fig. S7). This is in contrast to what has been reported for larger polyenes, such as nystatin, which causes formation of tension pores in response to osmotic stress, leading to accelerated cell death ^62^. The slightly protective effect of natamycin to osmotic pressure challenges could be due to an accelerated membrane permeation of the osmolyte glycerol, thereby lowering the osmotic pressure difference. Supporting that notion, we did not find evidence for osmotically induced water flow across the PM in the presence of natamycin using electron spin resonance (ESR) spectroscopy of the probe TEMPO (not shown). This is in accordance with earlier results on other fungal species ^63^. Together, our results suggest that binding of natamycin to the yeast PM causes membrane stress resulting in enhanced permeation of PI and other small molecules. Future studies are warranted to uncover the exact molecular basis for this increased permeability.

Using intrinsically fluorescent analogs of ergosterol, DHE, and cholesterol, CTL, which are only minimally modified compared to their parent molecule, we show here for the first time that sterols have a rather homogeneous lateral distribution in the yeast PM. This is in contrast to the observations made by Grossmann and co-workers using the polyene macrolide filipin ^2^. Filipin is a polyene very similar to natamycin, and as we show here for natamycin, filipin is known to form aggregates in membranes ^68–70^. Thus, a clustered filipin distribution is not evidence for the existence of sterol domains but rather an intrinsic property of the polyene, once it is associated with the cell membrane. Co-localization of filipin with eisosome markers was solely judged from single line profiles in the study by Grossmann et al. (2007) ^2^, and we show here for natamycin, that a thorough statistical analysis of line profiles from many cells is needed to assess polyene clustering in different membrane compartments. Indeed, we found that natamycin clusters form inside and outside of eisosomes and that the extent of co-clustering varies widely (Fig. 4 and S4). The eisosome marker Sur7 has been shown to be essential for virulence of yeast, e.g., of *C. albicans* ^67^. Our finding that natamycin aggregates inside and outside of Sur7-YPet-containing eisosomes, make it unlikely that natamycin targets Sur7-controlled eisosome integrity.

Also, others could not confirm a preferred localization of filipin to eisosomes but found a rather homogenous filipin distribution in the yeast PM supporting that staining patterns can vary significantly ^51^. In support of our observations on a homogenous lateral distribution of DHE and CTL in the PM, we found in recent membrane fractionation studies that different yeast membrane compartments contain similar amounts of ergosterol despite widely varying protein composition ^8^. Our results do not exclude the possibility that the nanoscale organization of ergosterol is different in various yeast membrane domains, but they show that sterol abundance is comparable in different PM compartments. Using label-free imaging of natamycin in giant unilamellar vesicles we demonstrated that it binds preferentially to the liquid-ordered (lo) lipid phase induced by ergosterol compared to the lo phase with cholesterol ^18^. We also showed that natamycin disrupts lipid packing in this phase allowing for increased passage of small polar molecules such as dithionite. Based on these findings and the results presented here, it is tempting to speculate that the entire yeast PM has properties resembling that of a tightly packed ergosterol-dependent lo phase. These properties of the yeast PM cause efficient recruitment of natamycin, which likely intercalates into the bilayer, forms aggregates, and weakens the interactions of ergosterol with phospho- and sphingolipids. These processes compromise membrane integrity resulting in enhanced passage of PI and likely other molecules, such as small metabolites. Together with its effect on ergosterol-dependent nutrient transporters this results in cell death caused by natamycin ^15,17^. In particular, sequestration of ergosterol by natamycin could increase membrane permeability, as both, cholesterol and ergosterol lower the permeation of small molecules across lipid bilayers under most conditions ^71^. Thus, membrane-embedded natamycin monomers or oligomers could bind sterols and thereby weaken phospholipid-sterol interactions in the membrane with the consequence of compromised permeability barrier function of the PM. A direct test of this model would require parallel imaging of natamycin and sterols, but this is not possible with our current tools since the fluorescence spectra of DHE and CTL overlap too strongly with those of natamycin to be able to separate them in the microscope (not shown).

Yeast sphingolipids are unique and different from those in mammalian cells, which is why they represent an important lipid class and potential drug target for antifungal therapies ^49^. We showed recently by NMR spectroscopy that natamycin weakens the interaction of stearoyl sphingomyelin (SSM) with ergosterol and interferes with phase separation in ternary mixtures of SSM, ergosterol, and palmitoyl-oleoyl phosphatidylcholine ^18^. Here, we show that inhibiting sphingolipid synthesis using myriocin increased binding of natamycin to yeast cells resulting in enhanced antifungal activity (Fig. 6). In contrast to model membranes, the yeast PM has an asymmetric lipid composition with the majority of ergosterol in the inner and likely most sphingolipids in the outer leaflet, causing a segregation of both lipid species in the PM ^10,11^. Due to their long alkyl chains intercalating into the inner leaflet, sphingolipids can impact interleaflet coupling and thereby affect the transbilayer sterol distribution, as shown in model membranes as well as yeast and mammalian cells ^10,72^. Supporting that model, we showed recently that myriocin treatment increases the sterol pool in the outer leaflet of the yeast PM ^10^. As a consequence of an increased ergosterol fraction in the outer PM leaflet, more natamycin can bind to yeast, resulting in more efficient killing of the cells. Based on these results we suggest that treatment of yeast with natamycin can be combined with inhibition of sphingolipid synthesis, which causes a redistribution of ergosterol in the PM and thereby enhanced recruitment and antifungal activity of the polyene. This might be a promising approach for future therapies of infections with pathogenic yeast species and could also involve other sphingolipid-specific antifungals and polyene macrolides.

## Materials and methods

### Reagents

The sterols ergosterol (45480), cholesterol (C8667), and DHE (810253P) including myriocin (M1177) and natamycin (32417) were purchased from MERCK KGaA. Cholestatrienol (CTL) was synthesized as described (see protocol and references in [33]). Propidium Iodide (P1304MP) and FM 4–64 (T3166) was purchased from Thermo Fisher.

### Yeast strains and culture conditions

The yeast strain hem1δ (*hem1*Δ cells) is derived from W303-1α (MATαade2-1 his3-11,15 leu2-3,112 trp1-1 ura3-1 can1-100) and was kindly provided by Dr. Thomas Pomorski (University of Copenhagen, Section Transport Biology). The yeast strain Sur7-Ypet and Pma1-YPet were cultured as described in ^7^. Yeast strain hem1δ were cultured in 5 ml media in a 50 ml Falcon tube at 30 ℃,150 rpm in YPD medium containing 2% (w/v) of glucose, 2% (w/v) of peptone, 1% (w/v) of yeast extract, and 200 mg/ml of adenine. The medium was supplemented with 20 μg/ml of *δ*-aminolevulinic acid (dALA) to ensure heme synthesis and thereby ergosterol biosynthesis when needed. Sur7-Ypet and Pma1-Ypet cells were grown under the same conditions as above but cultured in SD media containing 2% (w/v) glucose, 0.7% (w/v) yeast nitrogen base (YNB w/o amino acid), 0.19% (w/v) drop-out medium without uracil, 0.012% (w/v) adenine, and 0.002% (w/v) uracil. Experiments testing the impact of hyperosmotic stress were carried out using mixtures of water and glycerol as described ^37,73^.

### Labeling of yeast cells for fluorescence imaging

Exogenous sterols, ergosterol and cholesterol or their fluorescent analogues, DHE and CTL, respectively, were introduced to yeast strain *hem1*Δ. For that, cells were grown in YPD media without dALA (final concentration of 0.1OD_600_/ml) and supplemented with sterols loaded on Tween80 (final concentration of 5 μg/ml) and incubated for 22h at 30 °C and 150 rpm. After incubation, the yeast cells were washed three times with PBS. Cells were diluted in PBS to a final OD_600_ of 1.0. A final concentration of 50 μg/ml natamycin from a stock solution in methanol was added to yeast for 2h unless noted otherwise. In order to inhibit sphingolipid synthesis cells were treated with myriocin (final concentration of 15 μg/ml) for 10 min and afterward diluted to a final concentration of 5 μg/ml before adding natamycin. For staining dying cells, propidium iodide (final concentration of 60 μg/ml) was added for 5 min before imaging. For imaging, yeast cells were placed on poly-D-lysine coated dishes and allowed to settle for 10 min before washed one time with PBS to remove non-adherent cells.

### Fluorescence microscopy

The fluorescence images were obtained from two different wide-field microscopes. UV-sensitive epifluorescence microscopy was done using a Leica DMIRBE microscope with a 63x 1.4 NA oil immersion or 100x 1.3 NA oil immersion Fluotar objective, respectively (Leica Lasertechnik GmbH). A Lambda SC smart shutter (Sutter Instrument Company) was used as illumination control. The microscope contains an 10x extra magnification lens in the emission light path, resulting in a final pixel size of 193 nm and 123 nm, for the 63x and 100x objective, respectively. Natamycin, DHE and CTL were imaged using a specially designed filter cube obtained from Chroma Technology (Corp., Brattleboro, VA, USA) with a 335-nm (20-nm bandpass) excitation filter, 365-nm dichromatic mirror, and 405-nm (40-nm bandpass) emission filter. YPet-Sur7 was imaged using a standard green filter set (470-nm (20-nm bandpass) excitation filter, 510-nm dichromatic mirror, and 537-nm (23-nm bandpass) emission filter), while PI was imaged using a standard red filter set consisting of an excitation filter (535-nm, (50-nm bandpass), a 565-nm long-pass dichromatic filter and a 610-nm (75-nm bandpass) emission filter. For DHE, CTL and natamycin bleach stacks with 100 frames were routinely recorded with 400 msec acquisition time. For time-lapse imaging, all three channels (i.e., UV, green and red channel) were acquired sequentially every 30 min of incubation. Between image acquisitions, the focus was adjusted to account for chromatic aberration between the UV channel on one hand, and the green and red channels on the other, as described ^74^.

For more automated imaging of PI penetration into cells, a Nikon Ti-2 Eclipse microscope with a 100 x 1.4 NA oil immersion objective and post-magnification was used. Images were obtained from 15 randomly chosen areas of the culture dish using the circular random selection function with a radius of 0.5 mm and perfect focus control. The standard TRITC filter cube was used to image PI.

### Soft X-ray tomography

Sur7-YPet yeast cells were brought alive to the electron storage ring BESSY II (Berlin, Germany). They were maintained in Falcon tubes containing 50 ml of the growth media. Upon arrival, they were, centrifuged, washed with PBS and incubated with natamycin (final conc. 50 μg/ml in PBS) for 2h at room temperature. In some experiments, cells were washed two times with PBS, while in others, the natamycin solution remained at the sample before blotting and plunge freezing. Cells were seeded on poly-D-lysine coated R2/2 grids (Quanti-foil, 100 Holy Carbon Films, Grids: HZB-2 Au), and a small volume of 270 nm gold beads were added as fiducial markers to the grids. The sample was plunge-frozen in liquid ethane and stored in liquid nitrogen before imaging. Before cryo-freezing, water around yeast cells was rapidly removed by blotting, which is needed to prevent excessive ice formation on the sample during SXT imaging. Imaging of the samples was done on the U41 TXM beamline at BESSY II with 25-nm zone plates over a tilt range of 120-125° with 1° tilt steps at 510 eV. The camera exposure time was 6-12 s and the image pixel size is 9.8 nm. For alignment of tomogram frames, B-soft was used, while the Tomo3D software was used for reconstructions based on the filtered back-projection algorithm ^75,76^.

### Image analysis for fluorescence microscopy data sets

#### Preprocessing and deconvolution

Autofluorescence correction is done for all UV images (natamycin, DHE and CTL) by subtracting the last image of each bleach stack from the first one in an automated manner using batch processing in ImageJ. The resulting images only contain the bleaching fluorescent probe signal and were subsequently deconvolved using the ImageJ plugin DeconvolutionLab ^77^. The Richard-Lucy algorithm was used with 30 iterations (otherwise stated in the Supplemental Results). A theoretical PSF was used for deconvolution and generated using the Diffraction PSF 3D plugin in ImageJ (https://imagej.net/plugins/diffraction-psf-3d). Settings were chosen according to the used channel and camera specifications.

#### Extraction of line profiles, cluster and correlation analysis

Deconvolved images of natamycin, DHE or CTL were cropped on selected cells and using the segmented line selection tool in ImageJ, intensity profiles of 3 pixels width were extracted along the PM. The same approach was used on aligned stacks of images for each channel. A Macro was made to automatically measure intensity profiles and distances in microns for each image in a stack. The extracted line profiles were imported into Jupyter notebooks for further analysis (https://jupyter.org/ject). For the cluster analysis, the coefficient of variation (CV), which is the standard deviation of the line intensity divided by the mean, was calculated for each data set. To determine, how variation of the data can affect the CV, simulations were carried out. For those, the line profile was considered as a 1-dimensional fluorescence intensity signal, F(x) as a function of pixel position (i.e., spatial dimension x in µm), which is simulated using a periodic function:

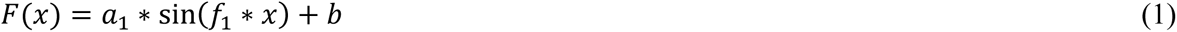

Here, x is the position along the line profile, the signal amplitude is *a*_1_, *f*_1_ is the frequency, and b is a background term, typically set to 200, which mimics the background signal in the fluorescence images. The signal is sampled at discrete positions x arranged in a Numpy array between 0.0 to 30.0 multiplied by the pixel size of the microscope camera ^78^. Additionally, the two-frequency equivalent of this function is used to simulate more complex intensity variations in the data:

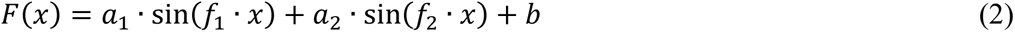

Peak detection was performed using routines in the Python library Scipy ^79^. The normalized line profiles (normalized to the maximum values) extracted from the natamycin fluorescence signal was used as a 1-D array to find all peaks using local maxima by comparison of neighbouring values using as search parameters height = 0.4 and width = 1. The parameters (distance, prominence, width, and threshold) are optimized to find the best detection of the peaks. Additionally, the peak widths are found and the relative half-maximal height for each peak.

For correlating the line profiles from the same cell but different markers, two approaches were used; first, the Pearson coefficient (PC) was calculated using the Python library SciPy ^79^. Second, a Bayesian analysis of the correlation was implemented based on the code provided in ^80^. For that, Monte Carlo sampling of the parameters and their posterior distribution is carried out using the NUTS sampler in PyMC3, a Python library for Bayesian inference of experimental data ^44^. The Bayesian analysis recovers not only the mean value of the estimated parameter but the entire PDF from which statistical measures, such as the highest density interval (HDI) can be extracted (see Results for details).

A Gaussian Mixture Model (GMM) was used for finding individual populations of the measured distribution of Pearson coefficients and CVs acquired for many cells. This was implemented using the Sklearn library in Python (https://scikit-learn.org/stable/). The method uses a maximum a posteriori estimation to find the most likely cluster assignment of the data for a given number of Gaussian distributions.

#### Segmentation of cells using cellpose

Images acquired from the Nikon Ti-2 Eclipse microscope were segmented using the deep learning-based segmentation method Cellpose, as described ^81^. The brightfield images were segmented using the built-in Cytoplasm 2.0 model (‘cyto2’) in Cellpose with a diameter of 85. For the fluorescent channel (PI) the background fluorescence was taken into account, and only cells with a mean intensity of 20 (after background correction) were denoted as dead. For all images, the total number of cells and the number of dead cells were counted. The entire process was automated by using a customized Python script.

## Acknowledgements

This research was funded by the Deutsche Forschungsgemeinschaft (DFG), MU 1017/12-1 (P.M.), WE1850-12/1 (P.W.) and D.W. as Mercator Fellow, as well as by NWO Graviation program BaSyC to BP. DW acknowledges funding from the Villum Foundation (grant no. 35865) and from the Danish Research Council (grant ID: 2032-00139B) as well as access to microscope equipment from the Danish Molecular Biomedical Imaging Center (DaMBIC, University of Southern Denmark).

---

**Figure S1.**
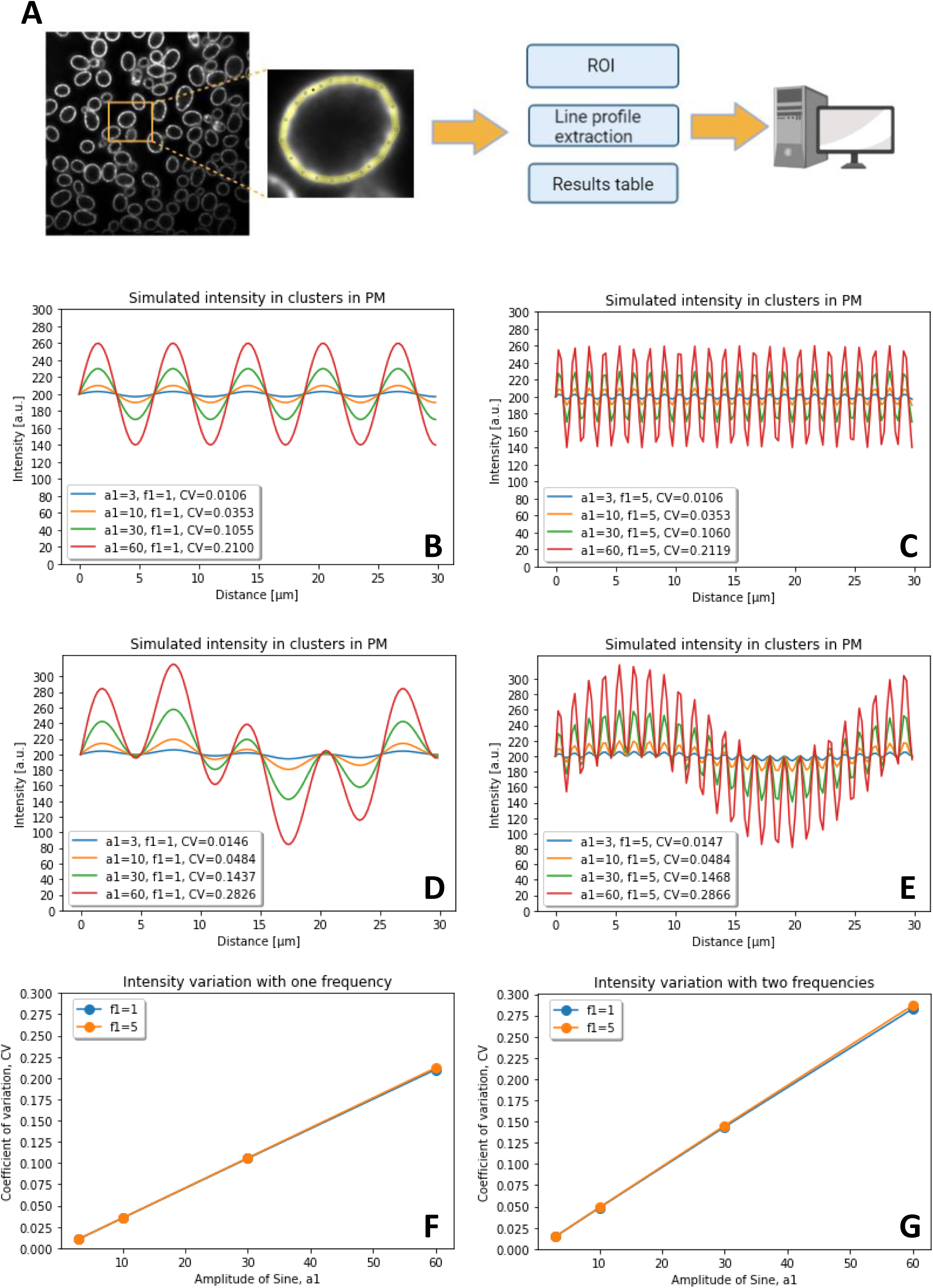
Procedure of extraction of line profiles and simulations using the coefficient of variation. Line profiles are drawn along the plasma membrane of yeast after deconvolution of the images as described in Materials and methods and shown in A. The obtained results table contains the measured positions along the membrane contour and the intensity profiles of the corresponding fluorescence channels (e.g., natamycin and Sur7-YPet). B-G, simulated intensity profiles based on Eq. 1 (B and C) or Eq. 2 (D and E) with differing amplitudes, *a*_1_ (B and D) or frequency, *f*_1_ (C and D). In D and E, the amplitude of the second component, *a*_2_, was identical to that of the first one (see Eq.2), while its frequency was lower (i.e., *f*_2_ = 0.25) to describe a slowly varying intensity component. The calculated coefficients of variation (CV) scale linearly with the change of amplitude and change with the number of oscillating components, but they do not vary much for different frequencies (F and G).

**Figure S2.**
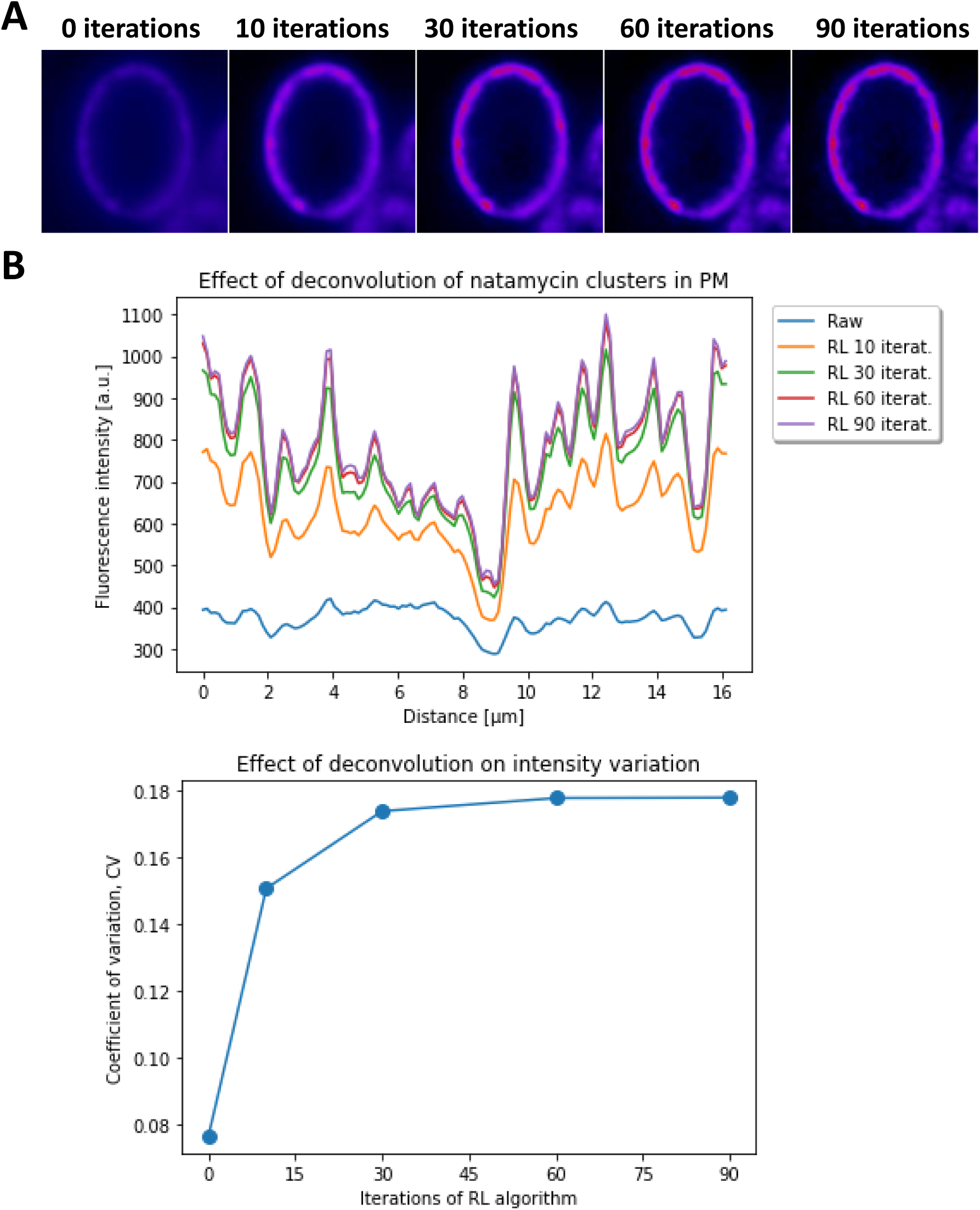
Effect of deconvolution of natamycin clusters in the PM of yeast. A, Deconvolution of a representative yeast cell using the Richard-Lucy algorithm in ImageJ as a function of iterations. B, extracted line profile from each iteration is plotted as fluorescence intensity against distance (µm). C, calculated coefficient of variation (CV), plotted as function of the number of iterations of the Richard-Lucy algorithm.

**Figure S3.**
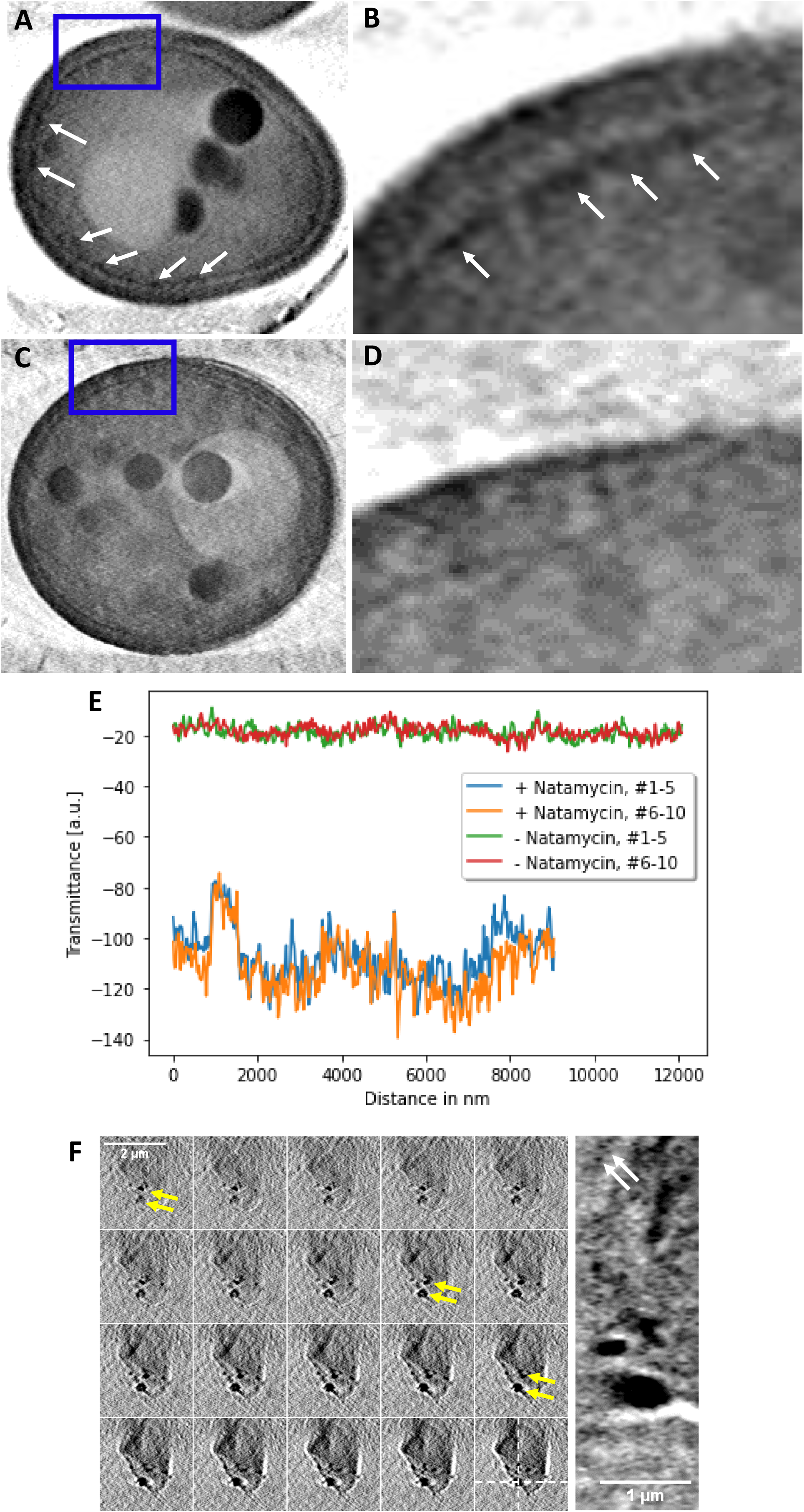
Soft X-ray tomography of natamycin aggregates. Yeast were either treated with natamycin for 2h (A and B) or left untreated (C and D) before plunge-freezing and tomographic imaging. A-D, sum projection of five central planes of a 3D reconstruction for natamycin-treated cells (A and B) or for untreated cells (C and D). B and D are zoomed versions of A and C at the respective blue box positions. Arrows in panel A and B point to putative natamycin aggregates. E, line profile extracted underneath the PM of natamycin-treated cells (blue and orange lines) or for untreated cells (green and red lines) for two subsequent sum projections of five slides each. F, 3D reconstruction of natamycin aggregates precipitated from buffer before plunge freezing. A montage of the central 30 frames of a large aggregate with about 2 µm in diameter having much smaller aggregates associated to it is shown in the left panel. These smaller aggregates are indicated by yellow arrows in selected frames. The right panel shows a a yz-view, i.e., along the optical axis of the reconstruction at the position indicated by the dashed white lines in the last frame of the montage. White arrows in the yz-view point to micro-aggregates with a diameter of about 100-250 nm, as they have been observed in cells.

**Figure S4.**
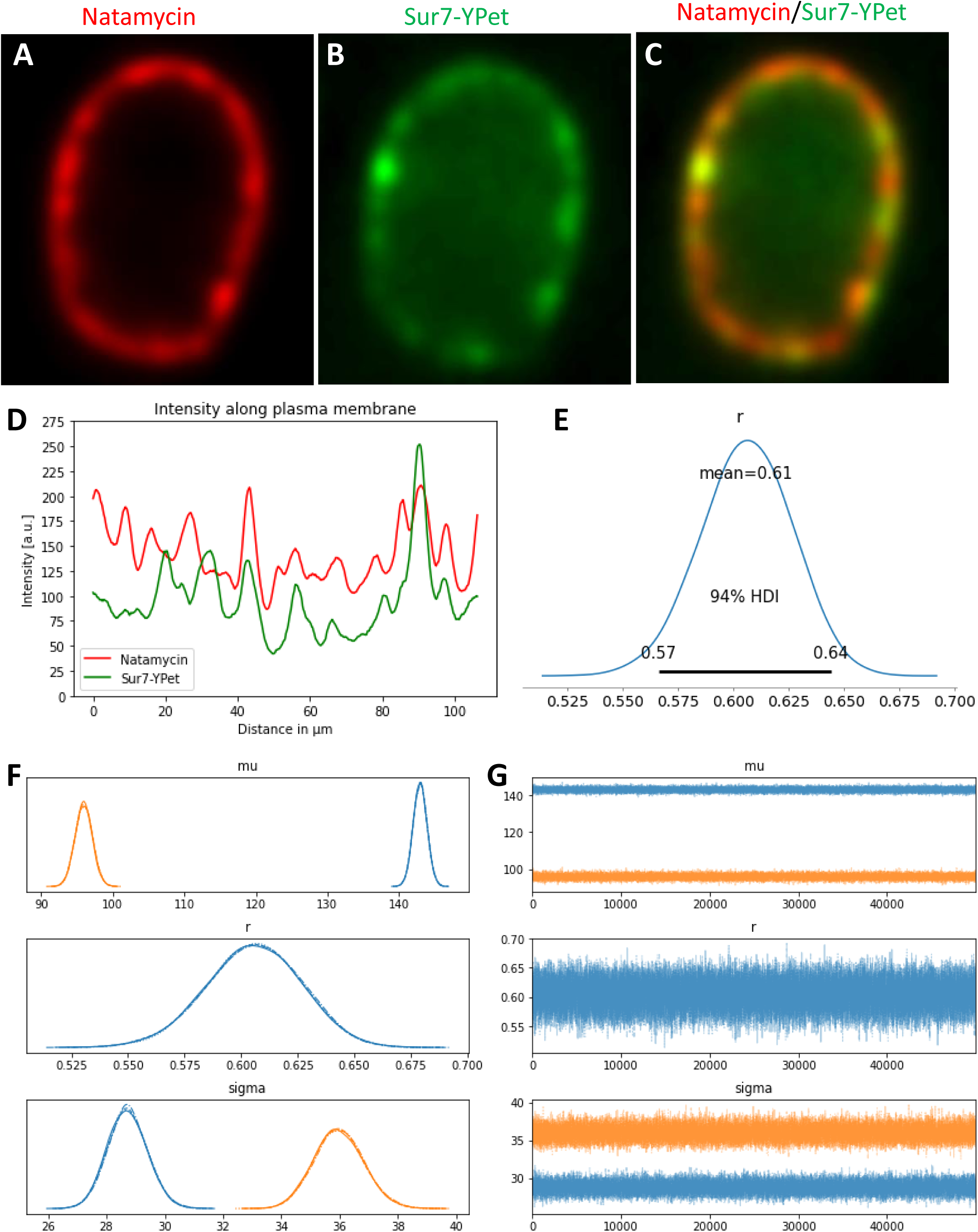
Example of natamycin clusters inside of eisosomes in the yeast PM. Yeast cells expressing Sur7-YPet, a green-fluorescent eisosome marker, were incubated with natamycin for 2h, washed and imaged. A, natamycin pseudo-colored in red, B, Sur7-YPet (green) and C, color overlay. D, line profiles for natamycin (blue line) and for Sur7-YPet (orange line) for the cell shown in A-C. E, probability density function (PDF) of Pearson coefficient, *r*, estimated by a Bayesian analysis with mean (r=0.52) and the highest density interval (HDI). F, Bayesian estimate of the PDF of mean intensity (upper panel) and variance (lowest panel) for each marker, i.e., posterior for Sur7-YPet (orange line) and natamycin (blue line) compared to the respective priors (dashed lines). The middle panel in F shows a comparison of posterior (straight line) with prior (dashed line) of the Pearson coefficient. G, corresponding MCMC runs as function of iteration. See main text and Figure 4 for further details.

**Figure S5.**
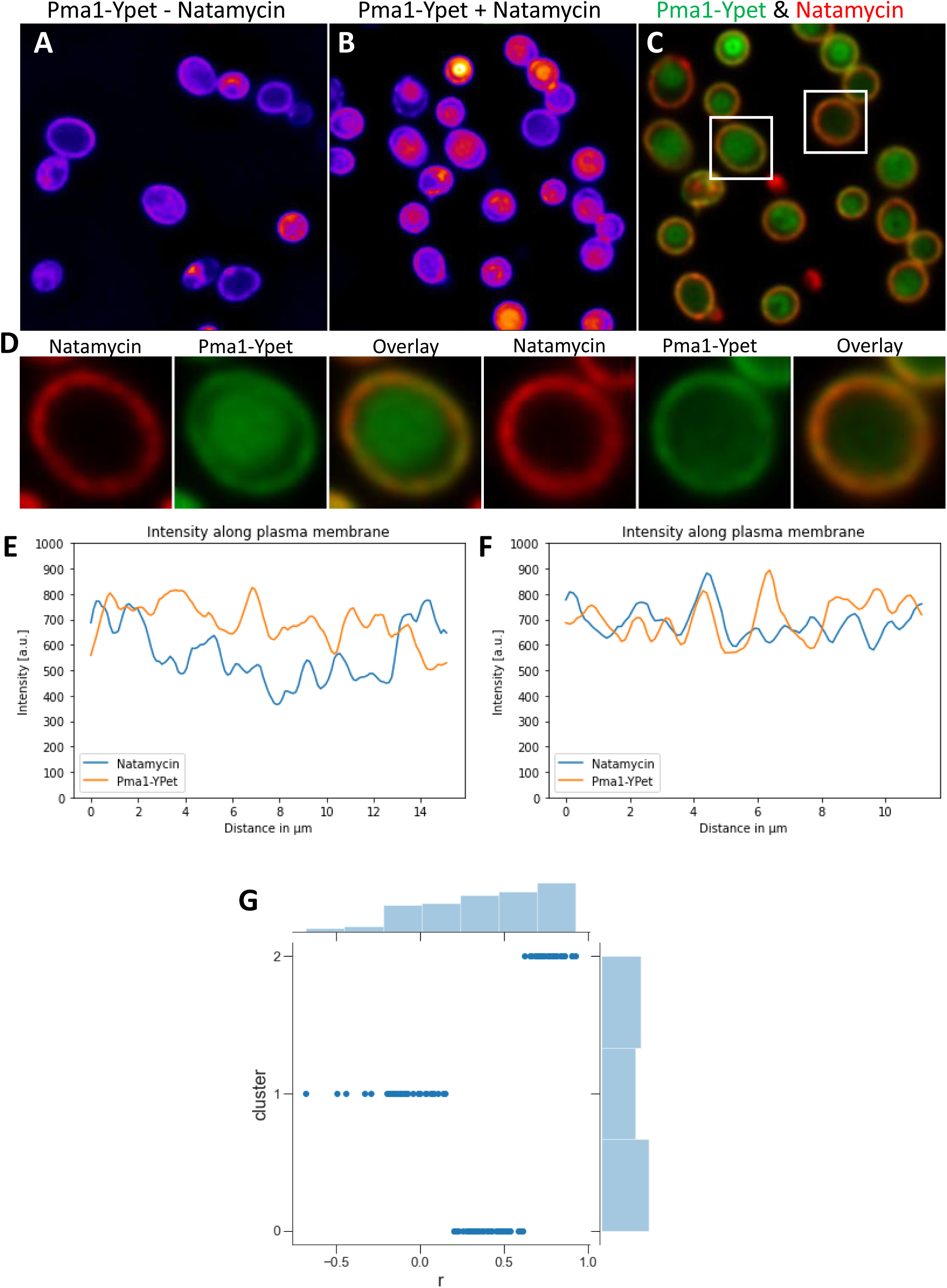
Co-localization of natamycin clusters with MCCs in the yeast PM. Yeast cells expressing Pma1-YPet, a green-fluorescent marker for MCCs, were incubated with natamycin for 2h, washed and imaged. A-C, Pma1-YPet in cells incubated without (A) or with 50 µg/ml natamycin (B) pseudo-colored with a FIRE-LUT to indicate intensities. C, color overlay with Pma1-YPet in green and natamycin in red. D, zoomed version of boxes shown in C with two representative cells with partial transport of Pma1-YPet to the vacuole (left panel) and most Pma1-YPet localized to the PM (right panel). E, line profiles were extracted from the PM of cells as shown for the left cell of D in E and for the right cell in D in F, respectively. Pearson coefficients were calculated for the entire population of analyzed cells (n=70) and deconvolved into three populations using a Gaussian Mixture model as shown in G.

**Figure S6.**
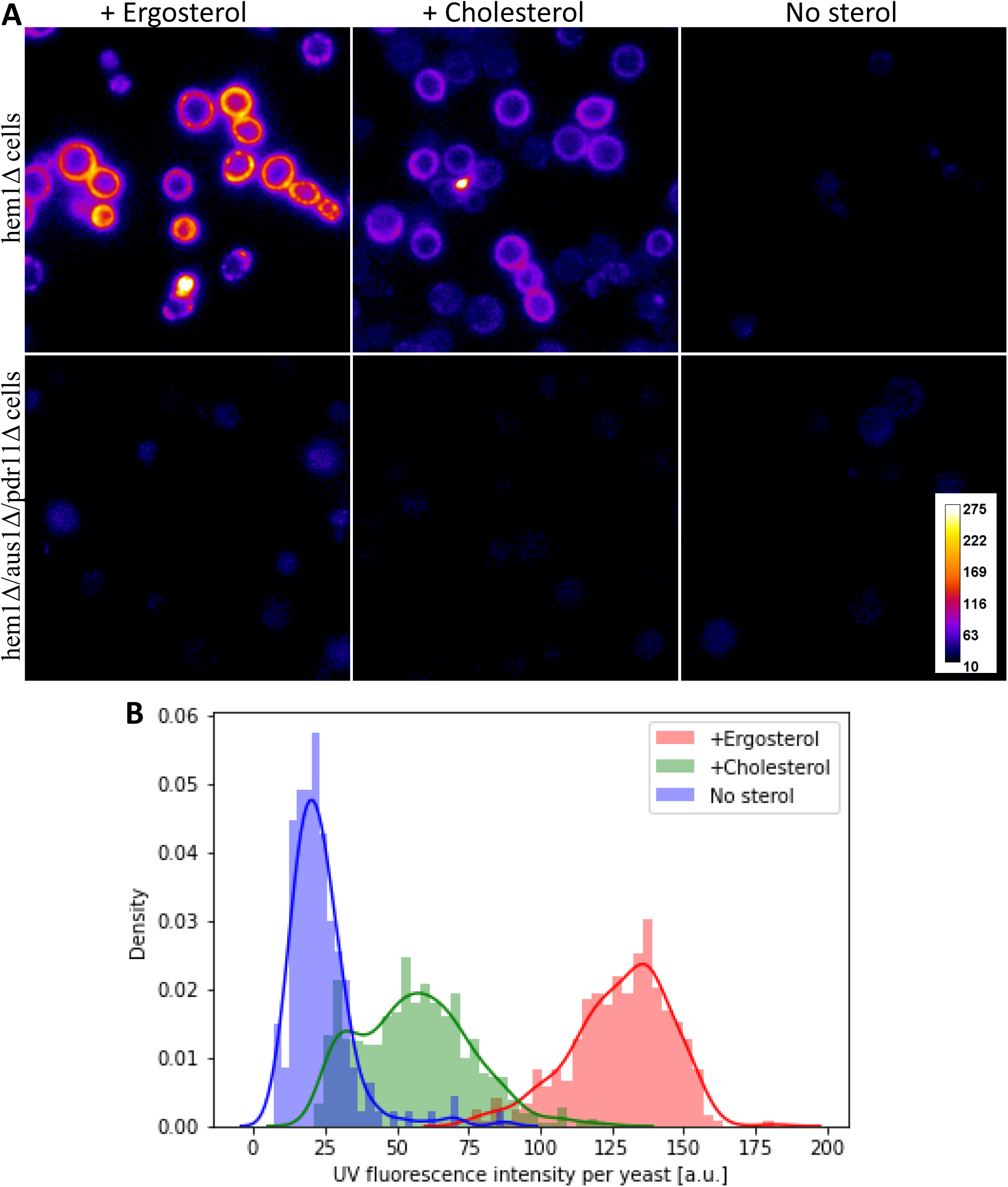
Binding of natamycin to yeast cells loaded with either ergosterol or cholesterol. Yeast cells with defective ergosterol synthesis (*hem1*Δ cells) or cells with an additional knock-down of the sterol importers Aus1/Pdr11 (i.e., *hem1*Δ*aus1*Δ*pdr11*Δ cells) were loaded with either ergosterol, cholesterol or no sterols at a concentration of 5 μg/ml in culture medium containing Tween80 for 22h at 30 °C. Cells were washed and incubated with natamycin (final concentration of 50 µg/ml) for 2h prior to imaging. A, representative images identically scaled and shown with a FIRE LUT with high intensities in yellow/orange, low intensities in dark blue and background in black. B, cell-associated fluorescence was quantified after thresholding the cells above background and plotted as kernel density plot for *hem1*Δ cells. Data is shown from one representative experiment with 5-8 image fields and ca. 50 cells per field for each condition.

**Figure S7.**
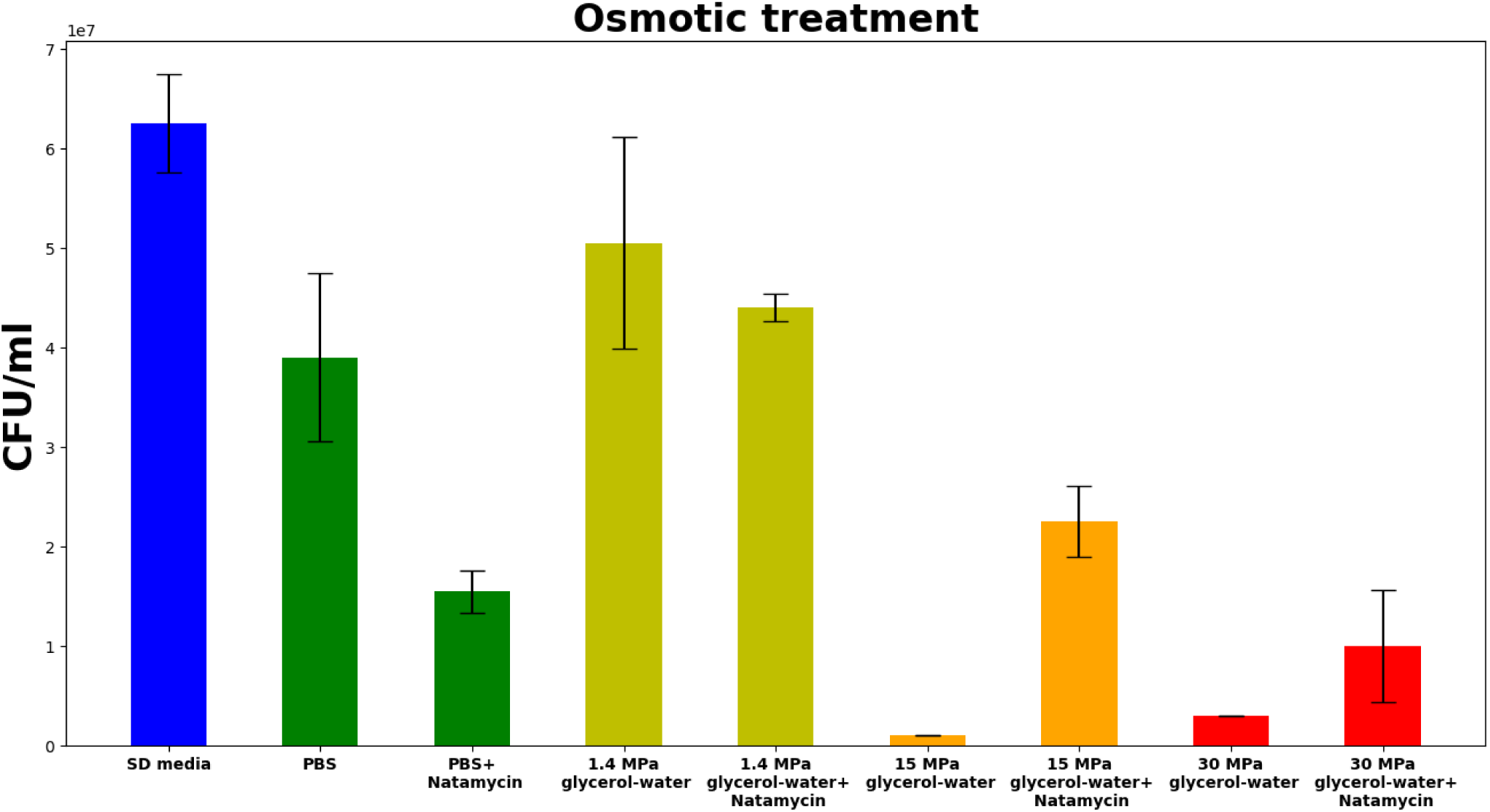
Natamycin-treated cells are not more sensitive to hyperosmotic stress. Sur7-Ypet cell were cultured under normal growth conditions in SD media. Cells were washed three times and diluted in solvents with various osmolarity corresponding to the indicated osmotic pressures to a final OD_600_ concentration of 0.5 OD/ml for 2h. Cells were plated on YPD plates and incubated at 30°C for 3 days before counting the colonies. Results are shown as mean +/-std of two technical replicates from one of two independent experiments.

## References

1 Ziolkowska, N. E., Christiano, R. & Walther, T. C. Organized living: formation mechanisms and functions of plasma membrane domains in yeast. Trends in cell biology 22, 151–158, doi:10.1016/j.tcb.2011.12.002 (2012).

2 Grossmann, G., Opekarova, M., Malinsky, J., Weig-Meckl, I. & Tanner, W. Membrane potential governs lateral segregation of plasma membrane proteins and lipids in yeast. The EMBO journal 26, 1–8, doi:10.1038/sj.emboj.7601466 (2007).

3 Stradalova, V. et al. Furrow-like invaginations of the yeast plasma membrane correspond to membrane compartment of Can1. Journal of cell science 122, 2887–2894, doi:10.1242/jcs.051227 (2009).

4 Gournas, C. et al. Conformation-dependent partitioning of yeast nutrient transporters into starvation-protective membrane domains. Proceedings of the National Academy of Sciences of the United States of America 115, E3145–E3154, doi:10.1073/pnas.1719462115 (2018).

5 Appadurai, D. et al. Plasma membrane tension regulates eisosome structure and function. Molecular biology of the cell 31, 287–303, doi:10.1091/mbc.E19-04-0218 (2020).

6 Malinska, K., Malinsky, J., Opekarova, M. & Tanner, W. Distribution of Can1p into stable domains reflects lateral protein segregation within the plasma membrane of living S. cerevisiae cells. Journal of cell science 117, 6031–6041, doi:10.1242/jcs.01493 (2004).

7 Bianchi, F. et al. Steric exclusion and protein conformation determine the localization of plasma membrane transporters. Nature communications 9, 501, doi:10.1038/s41467-018-02864-2 (2018).

8 van ’t Klooster, J. S., et al. Periprotein lipidomes of Saccharomyces cerevisiae provide a flexible environment for conformational changes of membrane proteins. eLife 9, doi:10.7554/eLife.57003 (2020).

9 Loura, L. M., Castanho, M. A., Fedorov, A. & Prieto, M. A photophysical study of the polyene antibiotic filipin. Self-aggregation and filipin--ergosterol interaction. Biochimica et biophysica acta 1510, 125–135, doi:10.1016/s0005-2736(00)00341-2 (2001).

10 Solanko, L. M. et al. Ergosterol is mainly located in the cytoplasmic leaflet of the yeast plasma membrane. Traffic 9, 198–214, doi:10.1111/tra.12545 (2018).

11 Aresta-Branco, F. et al. Gel domains in the plasma membrane of Saccharomyces cerevisiae: highly ordered, ergosterol-free, and sphingolipid-enriched lipid rafts. The Journal of biological chemistry 286, 5043–5054, doi:10.1074/jbc.M110.154435 (2011).

12 Guo, X. et al. Sterol Sponge Mechanism Is Conserved for Glycosylated Polyene Macrolides. ACS Cent Sci 7, 781–791, doi:10.1021/acscentsci.1c00148 (2021).

13 Hou, J., Daniels, P. N. & Burke, M. D. Small Molecule Channels Harness Membrane Potential to Concentrate Potassium in trk1Deltatrk2Delta Yeast. ACS chemical biology 15, 1575–1580, doi:10.1021/acschembio.0c00180 (2020).

14 Meena, M., Prajapati, P., Ravichandran, C. & Sehrawat, R. Natamycin: a natural preservative for food applications-a review. Food Sci Biotechnol 30, 1481–1496, doi:10.1007/s10068-021-00981-1 (2021).

15 te Welscher, Y. M., van Leeuwen, M. R., de Kruijff, B., Dijksterhuis, J. & Breukink, E. Polyene antibiotic that inhibits membrane transport proteins. Proceedings of the National Academy of Sciences of the United States of America 109, 11156–11159, doi:10.1073/pnas.1203375109 (2012).

16 te Welscher, Y. M., et al. Natamycin blocks fungal growth by binding specifically to ergosterol without permeabilizing the membrane. The Journal of biological chemistry 283, 6393–6401, doi:10.1074/jbc.M707821200 (2008).

17 Szomek, M. et al. Natamycin sequesters ergosterol and interferes with substrate transport by the lysine transporter Lyp1 from yeast. Biochim Biophys Acta Biomembr, 184012, doi:10.1016/j.bbamem.2022.184012 (2022).

18 Akkerman, V. et al. Natamycin interferes with ergosterol-dependent lipid phases in model membranes. Biochim Biophys Acta Advances 4, 100102 (2023).

19 Coutinho, A. & Prieto, M. Self-association of the polyene antibiotic nystatin in dipalmitoylphosphatidylcholine vesicles: a time-resolved fluorescence study. Biophysical journal 69, 2541–2557, doi:10.1016/S0006-3495(95)80125-6 (1995).

20 Aramwit, P., Yu, B. G., Lavasanifar, A., Samuel, J. & Kwon, G. S. The effect of serum albumin on the aggregation state and toxicity of amphotericin B. J Pharm Sci 89, 1589–1593, doi:10.1002/1520-6017(200012)89:12<1589::aid-jps10>3.0.co;2-6 (2000).

21 Fang, S. et al. Enhancing Water Solubility and Stability of Natamycin by Molecular Encapsulation in Methyl-β-Cyclodextrin and its Mechanisms by Molecular Dynamics Simulations. Food Biophysics 15, 188–195 (2020).

22 Qi, Z., Kang, Q., Jiang, C., Han, M. & Bai, L. Engineered biosynthesis of pimaricin derivatives with improved antifungal activity and reduced cytotoxicity. Appl Microbiol Biotechnol 99, 6745–6752, doi:10.1007/s00253-015-6635-9 (2015).

23 Strom, R., Crifo, C., Eusebi, F., Sabetta, F. & Oratore, A. Stoichiometry of hemolysis by the polyene antibiotic lucensomycin. Biochimica et biophysica acta 455, 961–972, doi:10.1016/0005-2736(76)90064-x (1976).

24 Tutaj, K. et al. Amphotericin B-silver hybrid nanoparticles: synthesis, properties and antifungal activity. Nanomedicine 12, 1095–1103, doi:10.1016/j.nano.2015.12.378 (2016).

25 Soo Hoo, L. Fungal fatal attraction: a mechanistic review on targeting liposomal amphotericin B (AmBisome((R))) to the fungal membrane. J Liposome Res 27, 180–185, doi:10.1080/08982104.2017.1360345 (2017).

26 Wasko, P. et al. Toward understanding of toxic side effects of a polyene antibiotic amphotericin B: fluorescence spectroscopy reveals widespread formation of the specific supramolecular structures of the drug. Mol Pharm 9, 1511–1520, doi:10.1021/mp300143n (2012).

27 Starzyk, J. et al. Self-association of amphotericin B: spontaneous formation of molecular structures responsible for the toxic side effects of the antibiotic. The journal of physical chemistry. B 118, 13821–13832, doi:10.1021/jp510245n (2014).

28 Gray, K. C. et al. Amphotericin primarily kills yeast by simply binding ergosterol. Proceedings of the National Academy of Sciences of the United States of America 109, 2234–2239, doi:10.1073/pnas.1117280109 (2012).

29 Anderson, T. M. et al. Amphotericin forms an extramembranous and fungicidal sterol sponge. Nat Chem Biol 10, 400–406, doi:10.1038/nchembio.1496 (2014).

30 Strom, R., Crifo, C. & Bozzi, A. The interaction of the polyene antibiotic lucensomycin with cholesterol in erythrocyte membranes and in model systems. I. A fluorometric and spectrophotometric study. Biophysical journal 13, 568–580, doi:10.1016/S0006-3495(73)86007-2 (1973).

31 Gruszecki, W. I., Gagos, M. & Herec, M. Dimers of polyene antibiotic amphotericin B detected by means of fluorescence spectroscopy: molecular organization in solution and in lipid membranes. J Photochem Photobiol B 69, 49–57, doi:10.1016/s1011-1344(02)00405-0 (2003).

32 Grudzinski, W., Sagan, J., Welc, R., Luchowski, R. & Gruszecki, W. I. Molecular organization, localization and orientation of antifungal antibiotic amphotericin B in a single lipid bilayer. Sci Rep 6, 32780, doi:10.1038/srep32780 (2016).

33 Lopes, S. & Castanho, M. A. R. B. Revealing the orientation of Nystatin and Amphotericin B in lipidic multilayers by UV-Vis linear dichroism. Journal of Physical Chemistry B 106, 7278–7282, doi:10.1021/jp020160s (2002).

34 Bolard, J. How do the polyene macrolide antibiotics affect the cellular membrane properties? Biochimica et biophysica acta 864, 257–304, doi:10.1016/0304-4157(86)90002-x (1986).

35 Szomek, M. et al. Direct observation of nystatin binding to the plasma membrane of living cells. Biochim Biophys Acta Biomembr 1863, 183528, doi:10.1016/j.bbamem.2020.183528 (2021).

36 Dupont, S., Beney, L., Ferreira, T. & Gervais, P. Nature of sterols affects plasma membrane behavior and yeast survival during dehydration. Biochimica et biophysica acta 1808, 1520–1528, doi:10.1016/j.bbamem.2010.11.012 (2011).

37 Dupont, S., Beney, L., Ritt, J. F., Lherminier, J. & Gervais, P. Lateral reorganization of plasma membrane is involved in the yeast resistance to severe dehydration. Biochimica et biophysica acta 1798, 975–985, doi:10.1016/j.bbamem.2010.01.015 (2010).

38 Legall, H. et al. Compact x-ray microscope for the water window based on a high brightness laser plasma source. Optics express 20, 18362–18369, doi:10.1364/OE.20.018362 (2012).

39 Larabell, C. A. & Le Gros, M. A. X-ray tomography generates 3-D reconstructions of the yeast, saccharomyces cerevisiae, at 60-nm resolution. Molecular biology of the cell 15, 957–962, doi:10.1091/mbc.e03-07-0522 (2004).

40 Uchida, M. et al. Quantitative analysis of yeast internal architecture using soft X-ray tomography. Yeast 28, 227–236, doi:10.1002/yea.1834 (2011).

41 Egebjerg, J. M., et al. Automated quantification of vacuole fusion and lipophagy in Saccharomyces cerevisiae from fluorescence and cryo-soft X-ray microscopy data using deep learning. Autophagy In press. (2023).

42 Juhl, A. D. et al. Niemann Pick C2 protein enables cholesterol transfer from endo-lysosomes to the plasma membrane for efflux by shedding of extracellular vesicles. Chem. Phys. Lipids 235, 105047, doi:10.1016/j.chemphyslip.2020.105047 (2021).

43 Juhl, A. D. et al. Quantitative imaging of membrane contact sites for sterol transfer between endo-lysosomes and mitochondria in living cells. Sci Rep 11, 8927, doi:10.1038/s41598-021-87876-7 (2021).

44 Salvatier, J., Salvatier, W., Thomas, V. & Fonnesbeck, C. Probabilistic programming in Python using PyMC3. PeerJ Computer Science 2, e55 (2016).

45 Solanko, L. M. et al. Ergosterol is mainly located in the cytoplasmic leaflet of the yeast plasma membrane. Traffic 19, 198–214, doi:10.1111/tra.12545 (2018).

46 Li, Y. & Prinz, W. A. ATP-binding cassette (ABC) transporters mediate nonvesicular, raft-modulated sterol movement from the plasma membrane to the endoplasmic reticulum. J. Biol. Chem. 279, 45226–45234 (2004).

47 Kohut, P. et al. The role of ABC proteins Aus1p and Pdr11p in the uptake of external sterols in yeast: Dehydroergosterol fluorescence study. Biochem. Biophys. Res. Commun. 404, 233–238 (2011).

48 Georgiev, A. G. et al. Osh proteins regulate membrane sterol organization but are not required for sterol movement between the ER and PM. Traffic 12, 1341–1355 (2011).

49 McEvoy, K., Normile, T. G. & Del Poeta, M. Antifungal Drug Development: Targeting the Fungal Sphingolipid Pathway. J Fungi (Basel) 6, doi:10.3390/jof6030142 (2020).

50 Grossmann, G. et al. Plasma membrane microdomains regulate turnover of transport proteins in yeast. The Journal of cell biology 183, 1075–1088, doi:10.1083/jcb.200806035 (2008).

51 Valdez-Taubas, J. & Pelham, H. R. Slow diffusion of proteins in the yeast plasma membrane allows polarity to be maintained by endocytic cycling. Current biology : CB 13, 1636–1640 (2003).

52 Wüstner, D. Plasma membrane sterol distribution resembles the surface topography of living cells. Mol. Biol. Cell 18, 211–228 (2007).

53 Adler, J., Shevchuk, A. I., Novak, P., Korchev, Y. E. & Parmryd, I. Plasma membrane topography and interpretation of single-particle tracks. Nat. Methods 7, 170–171 (2010).

54 Zumbuehl, A. et al. An amphotericin B-fluorescein conjugate as a powerful probe for biochemical studies of the membrane. Angewandte Chemie 43, 5181–5185, doi:10.1002/anie.200460489 (2004).

55 Silva, L., Coutinho, A., Fedorov, A. & Prieto, M. Conformation and self-assembly of a nystatin nitrobenzoxadiazole derivative in lipid membranes. Biochimica et biophysica acta 1617, 69–79, doi:10.1016/j.bbamem.2003.09.004 (2003).

56 Dos Santos, A. G., et al. The molecular mechanism of Nystatin action is dependent on the membrane biophysical properties and lipid composition. Physical chemistry chemical physics : PCCP 19, 30078–30088, doi:10.1039/c7cp05353c (2017).

57 Gonzalez-Damian, J. & Ortega-Blake, I. Effect of membrane structure on the action of polyenes II: nystatin activity along the phase diagram of ergosterol-and cholesterol-containing POPC membranes. J Membr Biol 237, 41–49, doi:10.1007/s00232-010-9301-2 (2010).

58 Recamier, K. S., Hernandez-Gomez, A., Gonzalez-Damian, J. & Ortega-Blake, I. Effect of membrane structure on the action of polyenes: I. Nystatin action in cholesterol- and ergosterol-containing membranes. J Membr Biol 237, 31–40, doi:10.1007/s00232-010-9304-z (2010).

59 Matsumori, N. et al. Direct interaction between amphotericin B and ergosterol in lipid bilayers as revealed by 2H NMR spectroscopy. Journal of the American Chemical Society 131, 11855–11860, doi:10.1021/ja9033473 (2009).

60 Silva, L., Coutinho, A., Fedorov, A. & Prieto, M. Nystatin-induced lipid vesicles permeabilization is strongly dependent on sterol structure. Biochimica et biophysica acta 1758, 452–459, doi:10.1016/j.bbamem.2006.03.008 (2006).

61 Nagi, M. et al. Serum cholesterol promotes the growth of Candida glabrata in the presence of fluconazole. Journal of infection and chemotherapy : official journal of the Japan Society of Chemotherapy 19, 138–143, doi:10.1007/s10156-012-0531-3 (2013).

62 Jokhadar, S. Z., Bozic, B., Kristanc, L. & Gomiscek, G. Osmotic Effects Induced by Pore-Forming Agent Nystatin: From Lipid Vesicles to the Cell. PloS one 11 (2016).

63 Van Leeuwen, M. R., Golovina, E. A. & Dijksterhuis, J. The polyene antimycotics nystatin and filipin disrupt the plasma membrane, whereas natamycin inhibits endocytosis in germinating conidia of Penicillium discolor. J Appl Microbiol 106, 1908–1918, doi:10.1111/j.1365-2672.2009.04165.x (2009).

64 Rizzo, J. et al. Coregulation of extracellular vesicle production and fluconazole susceptibility in Cryptococcus neoformans. mBio 14, e0087023, doi:10.1128/mbio.00870-23 (2023).

65 Grela, E. et al. Mechanism of Binding of Antifungal Antibiotic Amphotericin B to Lipid Membranes: An Insight from Combined Single-Membrane Imaging, Microspectroscopy, and Molecular Dynamics. Mol Pharm 15, 4202–4213, doi:10.1021/acs.molpharmaceut.8b00572 (2018).

66 Dong, P. T. et al. Polarization-sensitive stimulated Raman scattering imaging resolves amphotericin B orientation in Candida membrane. Sci Adv 7, doi:10.1126/sciadv.abd5230 (2021).

67 Douglas, L. M., Wang, H. X., Keppler-Ross, S., Dean, N. & Konopka, J. B. Sur7 promotes plasma membrane organization and is needed for resistance to stressful conditions and to the invasive growth and virulence of Candida albicans. mBio 3, doi:10.1128/mBio.00254-11 (2012).

68 Bridgman, P. C. & Nakajima, Y. Distribution of filipin-sterol complexes on cultured muscle cells: cell-substratum contact areas associated with acetylcholine clusters. J. Cell Biol. 96, 363–372 (1983).

69 Orci, L. et al. Heterogeneous distribution of filipin--cholesterol complexes across the cisternae of the Golgi apparatus. Proceedings of the National Academy of Sciences of the United States of America 78, 293–297 (1981).

70 McGookey, D. J., Fagerberg, K. & Anderson, R. G. Filipin-cholesterol complexes form in uncoated vesicle membrane derived from coated vesicles during receptor-mediated endocytosis of low density lipoprotein. J. Cell Biol. 96, 1273–1278 (1983).

71 Frallicciardi, J., Melcr, J., Siginou, P., Marrink, S. J. & Poolman, B. Membrane thickness, lipid phase and sterol type are determining factors in the permeability of membranes to small solutes. Nat. Comm. In press. (2022).

72 Courtney, K. C. et al. C24 Sphingolipids Govern the Transbilayer Asymmetry of Cholesterol and Lateral Organization of Model and Live-Cell Plasma Membranes. Cell reports 24, 1037–1049, doi:10.1016/j.celrep.2018.06.104 (2018).

73 Tran, K. & Green, E. M. Assessing Yeast Cell Survival Following Hydrogen Peroxide Exposure. Bio Protoc 9, doi:10.21769/BioProtoc.3149 (2019).

74 Wüstner, D. & Færgeman, N. J. Chromatic aberration correction and deconvolution for UV sensitive imaging of fluorescent sterols in cytoplasmic lipid droplets. Cytometry. Part A : the journal of the International Society for Analytical Cytology 73, 727–744 (2008).

75 Heymann, J. B. & Belnap, D. M. Bsoft: image processing and molecular modeling for electron microscopy. Journal of structural biology 157, 3–18, doi:10.1016/j.jsb.2006.06.006 (2007).

76 Agulleiro, J. I. & Fernandez, J. J. Tomo3D 2.0--exploitation of advanced vector extensions (AVX) for 3D reconstruction. Journal of structural biology 189, 147–152, doi:10.1016/j.jsb.2014.11.009 (2015).

77 Sage, D. et al. DeconvolutionLab2: An open-source software for deconvolution microscopy. Methods 115, 28–41, doi:10.1016/j.ymeth.2016.12.015 (2017).

78 Harris, C. R. et al. Array programming with NumPy. Nature 585, 357–362, doi:10.1038/s41586-020-2649-2 (2020).

79 Virtanen, P. et al. SciPy 1.0: fundamental algorithms for scientific computing in Python. Nature methods 17, 261–272, doi:10.1038/s41592-019-0686-2 (2020).

80 Singer, P. Bayesian Correlation with PyMC3, <https://medium.com/@ph_singer/bayesian-correlation-with-pymc-5dc6403e0599> (2015).

81 Stringer, C., Wang, T., Michaelos, M. & Pachitariu, M. Cellpose: a generalist algorithm for cellular segmentation. Nature methods 18, 100–106, doi:10.1038/s41592-020-01018-x (2021).

